# Co-evolution between PRDM9 and target sites and the Recombination Hotspot Paradox

**DOI:** 10.1101/2022.09.29.510084

**Authors:** Francisco Úbeda, Frederic Fyon, Reinhard Bürger

## Abstract

Recombination often concentrates in small regions called recombination hotspots where recombination is much higher than the genome’s average. In many vertebrates, including humans, gene PRDM9 specifies which DNA motifs will be the target for breaks that initiate recombination ultimately determining the location of recombination hotspots. Because the sequence that breaks (allowing recombination) is converted into the sequence that does not break (preventing recombination), the latter sequence is over-transmitted to future generations and recombination hotspots are self-destructive. Given their self-destructive nature, recombination hotspots should eventually become extinct in genomes they are observed. While empirical evidence shows that individual hotspots do become inactive over time (die), hotspots are abundant in many vertebrates: a contradiction called the Recombination Hotspot Paradox. What saves recombination hotspots from their foretold extinction? Here we formulate a co-evolutionary model of the interaction among sequence specific gene conversion, fertility selection and recurrent mutation. We find that when fertility selection is weaker than gene conversion, fertility selection cannot stop individual hotspots from dying but can save them from extinction by driving their re-activation (resuscitation). It can also save them from extinction by driving the birth of new hotspots in target sites with small allelic variation. The amount of allelic variation that can result in the birth of a hotspot depends on the strength of fertility selection and the mutation rate. In our model mutations balance death and resuscitation of hotspots maintaining their numbers over time. Interestingly we find that mutations are responsible for the oscillation of individual hotspots being asynchronous across the genome such that the average recombination across the genome remains constant. Our model thus contributes to better understanding how new hotspots may be formed thus explaining the Recombination Hotspots Paradox. From a more applied perspective our work provides testable predictions regarding the relation between mutation and fertility with life expectancy of hotspots.

## Introduction

Recombination —the exchange of genetic material between parental chromosomes— is a fundamental biological process that is key in generating DNA variability and trading information across genomes [22, 1, 42, 32]. It has a critical impact in ecology and evolution (selfish genes [17], genomic architecture [18], sexual reproduction [28], speciation [7] and conservation [13]), and in human disease (fertility [27, 36], cancer [15], association between genetic variants and illness [41, 40]). The impact of recombination relies not only on the intensity of recombination but also on its distribution across the genome. Contrary to prior beliefs, recombination is not uniformly distributed across the genome but concentrated in small chromosomal regions where recombination is ten to a thousand times more frequent than the genome’s average; these regions are known as *recombination hotspots* [33, 3, 30, 5]. In humans, most mammals and many vertebrates, the location of recombination hotspots is determined by the preference of alleles in PRDM9 locus for binding specific DNA motifs [26, 25, 4, 9]. Henceforth we will use the term recombination hotspots to refer to these PRDM9-directed recombination hotspots.

Empirical work shows that recombination is initiated by a double-strand break [29, 16, 33] and results in the conversion of the allele that breaks (*enabling recombination*) into the allele that does not break (*disabling recombination*) [33, 19, 20, 5]. The mechanism that initiates recombination provides an advantage in transmission to the allele disabling recombination over the allele enabling recombination [33, 19, 20, 5]. It is thus a matter of time that recombination hotspots become chromosomal regions where recombination is as frequent as the average across the genome [6] (henceforth *recombination coldspots).* This loss of activity is often referred to as the *death* of individual hotspots [10]. The self-destructive nature of recombination hotspots means that they should become extinct in genomes where they are initially found [6]. However, evidence shows that far from being rare, recombination hotspots are abundant [24, 3, 30, 5]. The *Recombination Hotspot Paradox* refers to the mismatch between the expected scarcity and observed abundance of recombination hotspots [6]: What saves recombination hotspots from their foretold extinction? Humans and chimpanzees do not share many recombination hotspots and even human subpopulations exhibit some level of recombination hotspot variation [38, 37, 48, 11, 43]. This suggests that the recombinational landscape is rapidly changing [38, 37, 48, 11, 43]. Furthermore, PRDM9 evolves extraordinarily rapidly to the point that has been considered the most rapidly evolving gene in many mammals [35]. Therefore a solution to the Recombination Hotspots Paradox that is consistent with empirical evidence should allow for (i) individual hotspots to die, (ii) the formation of new hotspots in sufficient numbers to compensate for the lost ones, and (iii) the limited number of hotspots shared by closely related species (rapidly changing recombinational landscape).

Because recombination underpins fundamental biological processes, the Recombination Hotspot Paradox has received much attention since it was formulated [2, 34, 8, 14, 31, 47, 21, 46]. In spite of the attention received, there is no fully satisfactory solution to the paradox. The initial attempts to solve the paradox explored whether the beneficial effects of recombination on fertility —in particular how recombination favours proper chromosomal segregation during meiosis thus preventing the formation of non-viable gametes [12, 27, 39, 36]— can save recombination hotspots from extinction [6, 34, 8, 31]. Mathematical models found that the allele that does not break at the recombination hotspot (disabling recombination) spreads in the population due to its transmission advantage, even when this allele reduces the fertility of the individual carrier [6, 34, 8, 31]. To maintain the allele enabling recombination, the benefits of recombination needed to be too strong to be realistic [6, 34, 8, 31]. Furthermore, when the benefits are strong enough, natural selection prevents individual hotspots from dying which is contrary to empirical observations on the death of individual hotspots and the rapidly evolving recombinational landscape [38, 37, 48, 11, 43, 35] (Figure **??**). These models, established that the paradox is well founded.

Advances in our understanding of the mechanisms initiating recombination found that, in many vertebrates, alleles at PRDM9 code for a protein that binds a specific sequence motif in hotspots [26, 25, 4]. Binding between protein and sequence causes a double-strand break (DSB) that initiates recombination at the binding site [25, 4]. This observation led to the verbal argument that mutations in PRDM9 could create new families of recombination hotspots that counteract the individual loss of recombination hotspots due to gene conversion [4]. The validity of this argument needed to be determined by a formal mathematical model. A first attempt to formalise this idea used an agent-based simulation as proof of principle that in small populations, drift, selection, conversion and mutation may lead to the formation of new recombination hotspots [47]. The computational power needed limited the size of the population simulated and the number of generations explored [47]. As a result of these limitation, it could not be established whether selection can sustain the birth of hotspots in the the long term or the genome would settle into intermediate levels of selection where hotspots are absent. In addition it could not evaluate how selection, conversion or mutation affects the turnover of recombination landscapes [21].

A second attempt to explore the evolution of PRDM9, formulated a one-locus stochastic model of the frequency of alleles at the PRDM9 locus when there is drift, selection, and mutation [21]. This work did not explicitly modelled the dynamics of alleles at target sites that bind PRDM9 [21]. To address this limitation the model assumed that mutations in PRDM9 result in fully functional recombination hostpots and that conversion in target sites were a function of the frequency of alleles at the PRDM9 locus —as opposed to the frequency of motifs at target sites [21]. However, the dynamics of alleles at PRDM9 can not be decoupled from the dynamics of alleles at its target sites; it is their co-evolution that determines whether target sites will be fully functional recombination hotspots or not and how conversion works as a function of the frequency of motifs at target sites [47, 46]. To fully understand the evolution of PRDM9 it is thus necessary to model the co-evolutionary dynamics between alleles at PRDM9 and alleles at their target sites. In addition this model implicitly assumes that there are potentially an infinite number of mutations in PRDM9 that produce new fully functional recombination hotspots without incurring in the fitness cost of starting recombination in motif in present in functional genes.

The most recent attempt to investigate the evolution of PRDM9, formulates a two-locus deterministic model of the co-evolutionary dynamics between PRDM9 and one of its target sites when there is selection and conversion [46]. This work found that when a hotspot has its binding motif replaced by a non-binding motif due to gene conversion, natural selection for the beneficial effects of recombination on fertility favours the spread of a mutant allele at PRDM9 that binds precisely the non-binding motif at the same recombination site [46]. This mathematical model shows that when the benefits of recombination are weak enough, natural selection favours the re-initiation of recombinational activity in the same chromosomal region where recombinational activity ceased due to conversion (*resuscitation* of a hotspot) [46]. However, this model finds that the life of a hotspot increases over time and eventually, in practical terms, hotspots no longer die [46]. This is contrary to empirical observations on the death of individual hotspots and the rapidly changing recombinational landscape [38, 37, 48, 11, 43, 35]. In addition, the ever increasing life of hotspots prevents the estimation of the expected lifetime of hotspots, the change in recombination landscape and how they scale with the selection and conversion.

Here we extend the most recent attempt to model the evolution of PRDM9 [46] by incorporating mutations both in PRDM9 loci and its target sites —an evolutionary force neglected in [46]. In addition, we extend [46] by considering multiple target sites. We thus formulate multilocus deterministic models of the co-evolutionary dynamics between PRDM9 and one or more of its target sites when there is selection, conversion and mutation. We find that mutations change significantly the co-evolutionary dynamics driving permanent and regular oscillations between high recombination and low recombination in the same target sites. Mutations thus prevent the system from getting stuck in hotspots or coldspots, a behaviour that proved to be problematic in previous models [46]. Unexpectedly, we find that mutations drive the alternation of target sites in their oscillation between high and low recombination. The outcome is that the average recombination across the genome remains constant while the recombination in each site oscillates widely. Finally, we extend our model to consider finite populations thus bridging the gap between our deterministic predictions and stochastic observations in pervious models [47]. From a theoretical perspective we discuss the implications of our model for solving the Recombination Hotspots Paradox. From a more applied perspective, we discuss the insights provided by our model into the mechanism and genomic distribution of PRDM9 directed recombination hotspots. These results allow us to predict how the expected lifetime of hotspots and the recombination landscape change with fertility, conversion, mutation (in each of the loci) and the number of targets thus opening the possibility of calibrating this model with empirical data.

## Methods

We model the interaction between one locus (the *modifier* locus) coding for a protein that recognizes specific motifs at one or more loci (the *target* loci) where crossover may be initiated. This is the behavior of gene PRDM9 controlling recombination hotspots in humans and many vertebrates [25, 4, 5, 9]. We start by presenting a deterministic two-allele version of the model with one modifier and one target locus. It describes the change in gamete frequencies under fertility selection, recombination, and mutation in a sufficiently large population. In Section 1 of the SM we introduce a generalised multi-allelic version from which we derive the simpler di-allelic case presented in the main text. Then we extend the deterministic model to two target loci, i.e., a three-locus two-allele model. Finally, we formulate a model assuming that the population is finite and that there may be any number of target loci. Throughout, following previous work we assume a randomly mating diploid population with non-overlapping generations [47, 21, 46].

### Deterministic one-target model

We model a modifier locus *A* that carries two alleles *A*_1_ or *A*_2_, each encoding a protein that attempts to bind a motif at a target locus *B*. The target locus *B* also carries two alleles *B*_1_ or *B*_2_, each corresponding to a base pair motif that the protein produced by locus A may attempt to bind. Let *x*_1_, *x*_2_, *x*_3_, *x*_4_ be the frequencies of the gametes *A*_1_*B*_1_, *A*_1_*B*_2_, *A*_2_*B*_1_, *A*_2_*B*_2_, respectively. We denote the recombination probability between modifier and target locus by *r_m_*. For the derivation of our results we assume 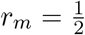, i.e., modifier and target locus are either far apart on the same chromosome or on separate chromosomes. This assumption is consistent with empirical observations and previous work on the topic [31, 46].

Random union of gametes results in diploid individuals whose genotype determines the outcome of meiosis in adults. We assume that alleles are equally likely to be expressed in each of the loci. As a result, each allele at the modifier locus is equally likely to produce the protein that will attempt to bind one of the two target motifs at random. We assume further that at most one binding attempt can occur per individual per generation during meiosis. We also assume that a match between subscripts of the alleles expressed at the modifier and target loci represents a match between recognition and target sequences. Thus a match between subscripts results in a DSB with probability *b* (where 0 < *b* < 1). A mismatch between subscripts yields no DSBs. Notice that two of the gametic types, *A*_1_*B*_1_ and *A*_2_*B*_2_,produce a protein that can bind the target (crossover enabling types), whereas the other two, *A*_1_*B*_2_ and *A*_2_*B*_1_, produce a protein that cannot bind the target (crossover disabling types).

During the DSB repair process there might be a crossover event in or near the target locus with probability *r_t_* (henceforth crossover probability) and none with probability 1 – *r_t_* (where 0 < *r_t_* < 1) [23, 33]. During the repair process the allelic motif that breaks is converted into the allelic motif that does not break with probability *c* and is restored to the allelic motif that breaks with probability 1 – *c* (where 0 < *c* < 1) [45, 44, 23, 33]. Notice that biased gene conversion results in the over-transmission of the allele that is less likely to break [6, 33].

Consistently with empirical observations and previous work, we assume that individuals undergoing crossover at the target locus have proper chromosomal segregation and do not suffer any fertility cost [12, 27, 39, 36, 6, 34, 31, 46]. Individuals that do not undergo crossover have defective chromosomal segregation producing non-viable gametes with probability *f* (where 0 < *f* < 1). Alleles at the modifier locus mutate from *A*_1_ to *A*_2_ or vice versa with probability *μ_A_*. Similarly, alleles at the target locus mutate from *B*_1_ to *B*_2_ or vice versa with probability *μ_B_*.

Gametic types segregate following Mendelian rules which brings us back to the beginning of the census. We write **x** = (*x_j_*) and 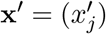 for the vectors that contain the four gametic frequencies in successive generations, *D* = *x*_1_*x*_4_ – *x*_2_*x*_3_ for the linkage disequilibrium, and **d** = (1, −1, −1, 1)^τ^ for the vector of signs of the linkage disequilibrium terms. We use ^τ^ for matrix or vector transposition. The 4 × 4 matrices **W** = (*w_ij_*) and **M** = (*m_ij_*) contain the fitnesses and the mutation probabilities, respectively, of each gametic type. The fitness matrix combines the effects of fertility selection and conversion:

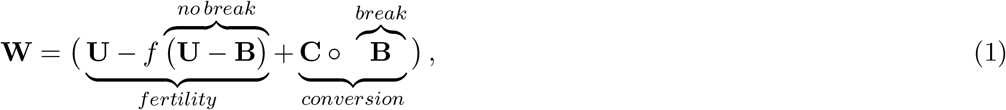

where **U** is a 4 × 4 matrix of ones,

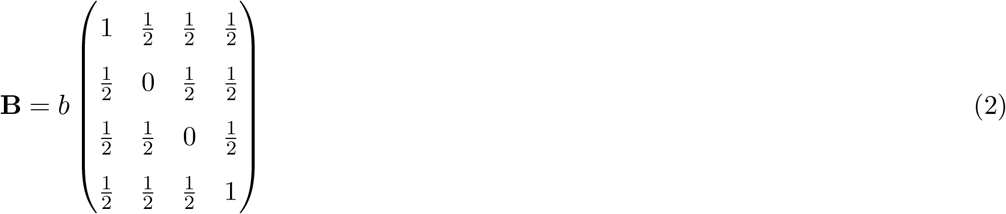

contains the break probabilities,

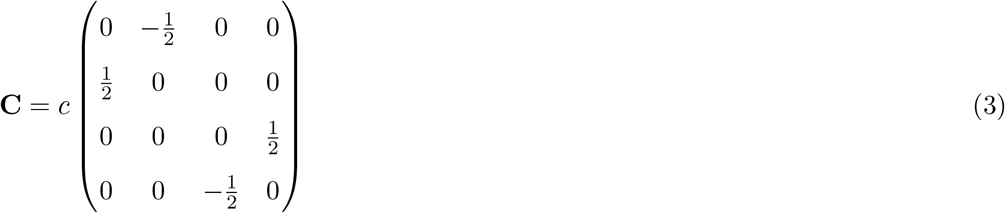

contains the conversion probabilities, and o denotes the pointwise (Schur) product of matrices. Finally,

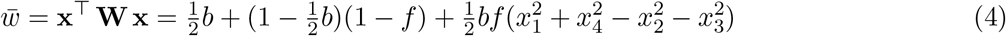

is the population mean fitness, which is not affected by conversion.

We note the mutation matrices at the two loci:

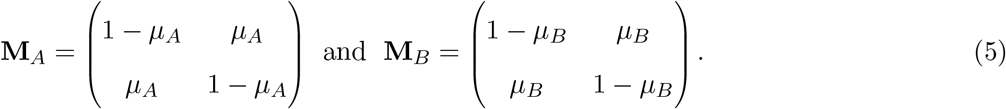

Because mutations occur independently, the resulting gametic mutation matrix **M** is their Kronecker product, i.e., **M** = M_*A*_ ⊗**M**B.

With these ingredients, the gametic frequencies after recombination, selection, and mutation are given by

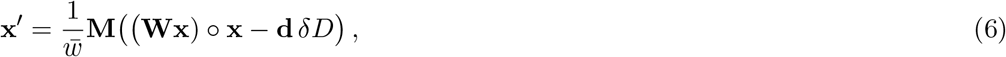

where *δ* is a function that measures the effect of fertility and conversion on linkage disequi librium:

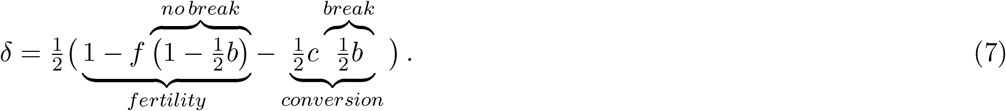

For detailed derivations, see Section 1 of the SM.

The phenotype we are interested in characterising is the population mean crossover probability at the target site:

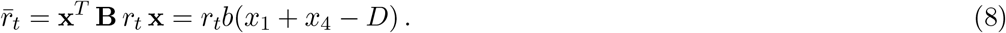

For simplicity, in this manuscript we assume *r_t_* = 1.

Sometimes, it is more insightful to consider the allelic frequencies and the linkage disequilibrium between them, instead of the gametic frequencies. Let *p* = *x*_1_+*x*_2_ and *q* = *x*_1_+*x*_3_ be the frequencies of alleles *A*_1_ and *B*_1_ respectively (see Section 2 of the SM for details on the transformation from (*x*_1_, *x*_2_, *x*_3_, *x*_4_) to (*p,q,D*)).

### Deterministic two-target two-allele model

In this section, we briefly outline a three-locus model that extends the previous two-locus model by adding a second locus (locus *C*) as potential target of the PRDM9 protein encoded by locus *A*. We assume that each locus can carry two alleles. We assume that during meiosis the PRDM9 protein attempts to bind one and only one target allele either at locus *B* or *C* with equal probability 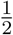. We continue to assume that a match between subscripts in locus *A* and *B* or *C* results in a DSB with probability *b*. Recurrent mutations happen in the three loci. All other interactions between the PRDM9 locus and its two target loci remain the same as in the previous section. Detailed equations can be found in Section 4 of the SM.

### Stochastic multi-target multi-allele model

In this section, we outline a stochastic version of the deterministic model introduced above to describe a populations with a finite number of individuals *N* that carry genomes with a variable number of target sites. We assume that target sites can carry the same pair of alleles (thus belonging to the same family) or a different pair of alleles (thus belonging to different families). PRDM9 can carry as many alleles as alleles segregate at all target sites while each target site only segregates 2 alleles. We still assume that there is only one binding attempt per meiotic event. We assume that a binding may be successful if and only if the subscripts of the alleles in the PRDM9 locus and the target match. Recurrent mutations happen in all loci considered. All other interactions between the PRDM9 locus and its target loci remain the same as in the original model. A detailed description of the code used in our simulations can be found in Section 5 of the SM. The code itself is available on GitHub, https://github.com/FredericFyon/Recombination-Hotspot-Unparadox.

## Results

Henceforth, for simplicity we assume that when there are recurrent mutations these occur in all (PRDM9 and target) loci at the same rate *μ* unless stated otherwise. When the mutation probability is small, we find two equilibria near the corners of the state space where the frequency of one of the two crossover enabling gametic types is high and the frequency of all other gametic types is low, **x**^*1^(close to fixation of *A*_1_*B*_1_) and **x**^*4^ (close to fixation of *A*_2_*B*_2_). We also find an internal equilibrium where all gametic types are present, **x**^*5^, and the allele frequencies *p* of *A*_1_ and *q* of *B*_1_ are 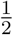 each. The internal equilibrium exhibits substantial linkage disequilibrium (see Section 3 of the SI for explicit expressions for these equilibria).

First consider the case when the transmission gain due to fertility selection *f* is stronger than the transmission gain due to gene conversion 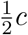, that is 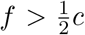. The two equilibria near the corners, **x**^*1^ and **x**^*4^, are within the state space with biological meaning. These equilibria are stable when the mutation probability is small (Figure 1; Section 3 of the SM). The internal equilibrium, **x**^*5^, is unstable when the mutation probability is small (Figure 1; Section 3 of the SM). The linkage disequilibrium at the internal equilibrium **x**^*5^ is positive, *D*\* > 0, thus crossover enabling gametic types are over-represented at the internal equilibrium. More specifically, stronger mutation weakens linkage disequilibrium.

**Figure 1:**
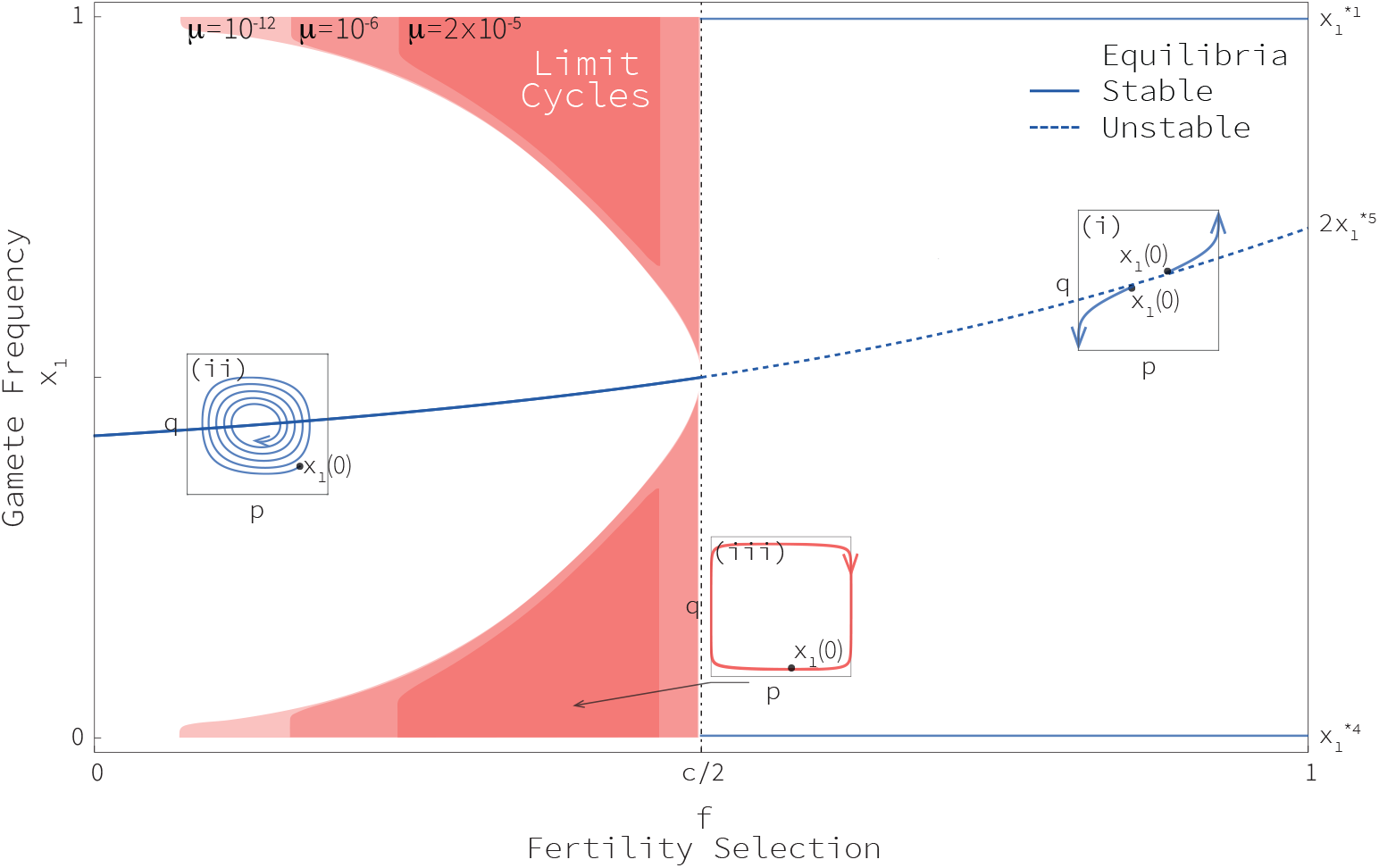
Basin of attraction of limit cycles and each equilibrium. This figure depicts the frequency of gametic type *A*_1_*B*_1_ (*x*_1_) as a function of the fitness cost *f* for given values of *r_m_,b,c,μ*. When 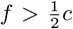 and *μ* > 0 equilibria **x**^*1^,**x**^*4^,**x**^*s^ are biologically meaningful. Equilibria **x**^*1^, **x**^*4^ are stable whereas equilibrium **x**^*5^ is unstable. The figure is constructed by starting trajectories at different initial conditions, sampled at equally spaced intervals along the line connecting corners (1, 0, 0, 0) and (0, 0, 0, 1) for which (*x*_1_(0), *x*_2_(0), *x*_3_(0), *x*_4_(0)) = (*x*_1_(0), 0, 0,1 – *x*_1_(0)). Initial conditions above the dotted curve (which represents equilibrium **x**^*5^) lead to equilibrium **x**^*1^ while those below **x**^*5^ lead to **x**^*4^. This behaviour is summarised using inset (i), which represents the dynamics in the allelic space (*p, q*). When 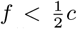 and *μ* > 0, only equilibrium **x**^*5^ is inside the region where *x*_1_ has biological meaning. Equilibrium **x**^*5^ is stable. Trajectories starting at initial condition *x*_1_(0) within the white region spiral towards the symmetric internal equilibrium (see inset ii). When initial conditions are within the red region, the solutions oscillate towards a limit cycle (see inset iii). As the mutation probability increases, the set of initial conditions converging to the stable limit cycle shrinks.

When 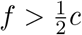 then the trajectories converge to one of the equilibria **x**^*1^ and **x**^*4^ and one of the crossover-enabling gametes gets nearly fixed (Figure 1). This recovers previous results indicating that when selection is unrealistically strong, it prevents the death of individual recombination hotspots and promotes a stable recombinational profile [6, 34]. Henceforth we will focus on the more realistic case when selection strength is weaker than conversion.

In the following, we consider the case when fertility selection is weaker than gene con-version, that is 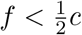. Then the two equilibria near the corners, **x**^*1^ and **x**^*4^, are outside of the state space with biological meaning (Figure 1; Section 3 of the SM). The internal equilibrium, **x**^*5^, is stable when the mutation probability considered is small (Figure 1; see Section 3 of the SM). The linkage disequilibrium at the internal equilibrium **x**^*5^ is negative, D^*^ < 0, thus crossover disabling gametic types are over-represented at the internal equilibrium. Again, stronger mutation weakens linkage disequilibrium, i.e., makes it less negative.

Numerical analysis shows that with fertility selection, recombination and mutation, trajectories either approach the internal equilibrium or approach a limit cycle depending on the initial conditions (Figure 1). We mapped the set of initial conditions leading to one dynamic behavior or the other using numerical methods (Figure 1). The greater the mutation probability the smaller the set of initial conditions leading to a limit cycle (Figure 1). When initially one of the gametic types is abundant and the others rare, the frequencies of the gametic types converge towards the limit cycle and oscillate regularly over time (Figure 2.A). Consistently with previous results, in the absence of mutations the amount of time spent in each of the corners where one gametic type is almost fixed increases in each oscillation [46] (Figure 2.B.i). Eventually the gametic frequency shows no perceptible change and the systems behaves, in practical terms, as if stuck in one of the corners (Figure 2.B.i). The introduction of mutations in the PRDM9 locus changes significantly this behavior (Figure 2.B.ii). With mutations the amount of time spent in each of the corners where one gametic type is fixed remains constant over time and the system can oscillate perpetually (Figure 2.B.ii).

**Figure 2:**
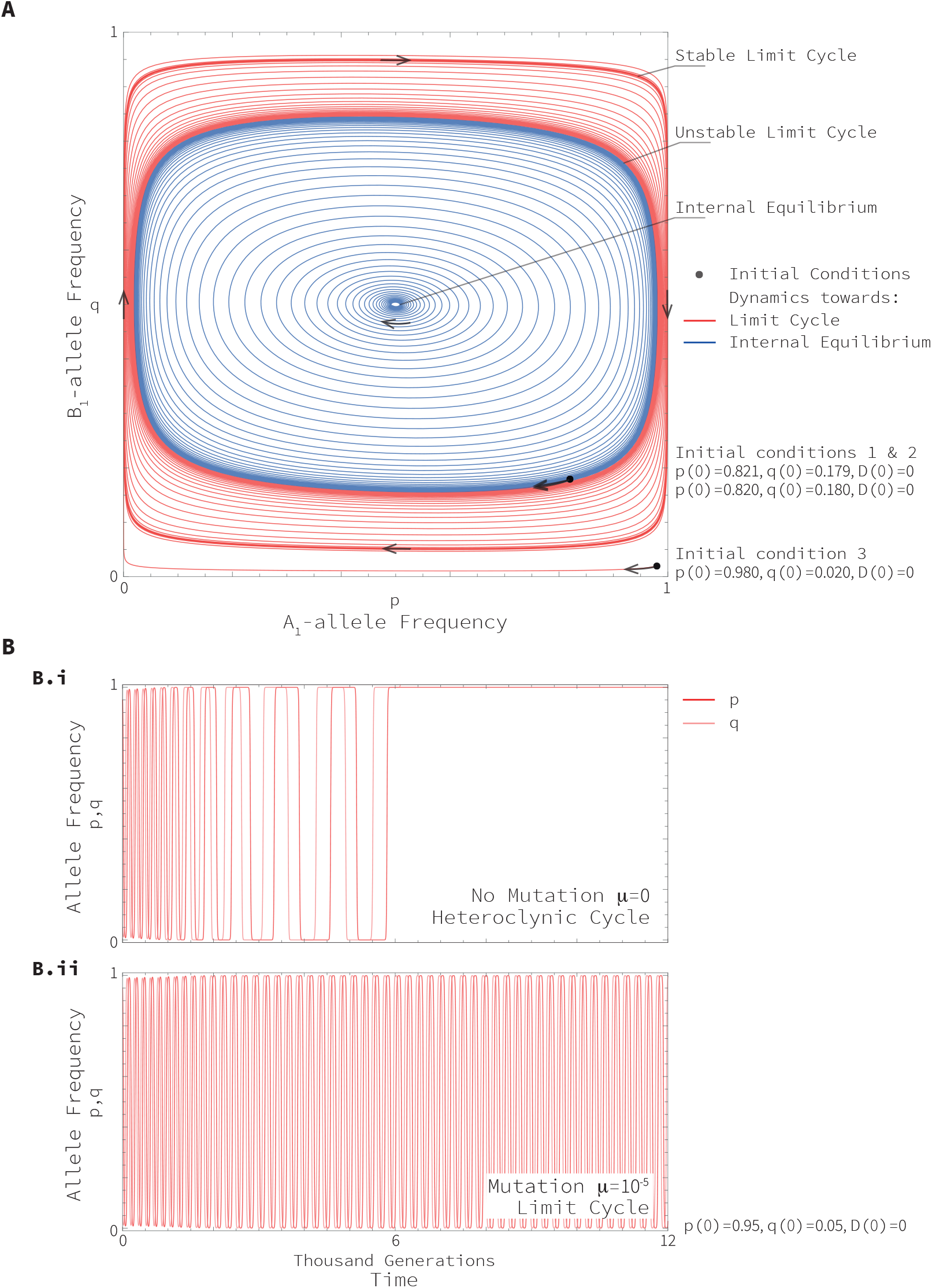
Examples of dynamics of the allelic frequencies in the parameter region 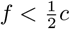. **Panel A** displays the dynamics in (p, q)-space. It shows the existence of a stable limit cycle, an internal equilibrium and an unstable limit cycle separating them. Red and blue are used to represent the trajectories leading to the stable limit cycle and the internal equilibrium respectively. The trajectories correspond to the set of parameter values *f* = 0.4, *b* = *c* =1, 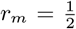, *μ* = 10^-5^. **Panel B** displays the frequencies of *p* and *q* as a function of time. **Subpanel B.i** provides an example when there is no mutation, *μ* = *μ_A_* = *μ_B_* = 0. **Subpanel B.ii** provides an example when there is mutation, *μ* = *μ_A_* = *μ_B_* = 10^-5^. Both trajectories correspond to the set of parameter values *f* = 0.26, *b* = 1, 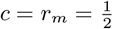.

The number of generations it takes for a cycle of gametic types to be completed (the *period* of the limit cycle) changes with the selection strength and the mutation probability. The closer the strength of fertility selection is to 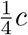, the longer the period of the limit cycle (Figure 3.A). The greater the mutation probability, the shorter the period of the limit cycle (Figure 3.A). The period of the limit cycle is relevant because measures, indirectly, the life expectancy of recombination hotspots. The greater the period of the limit cycle the greater the life expectancy of recombination hotspots (see SM). In the absence of mutation the period increases from one oscillation to the next and it is not possible to calculate neither a period nor a life expectancy for the recombination hotspots.

**Figure 3:**
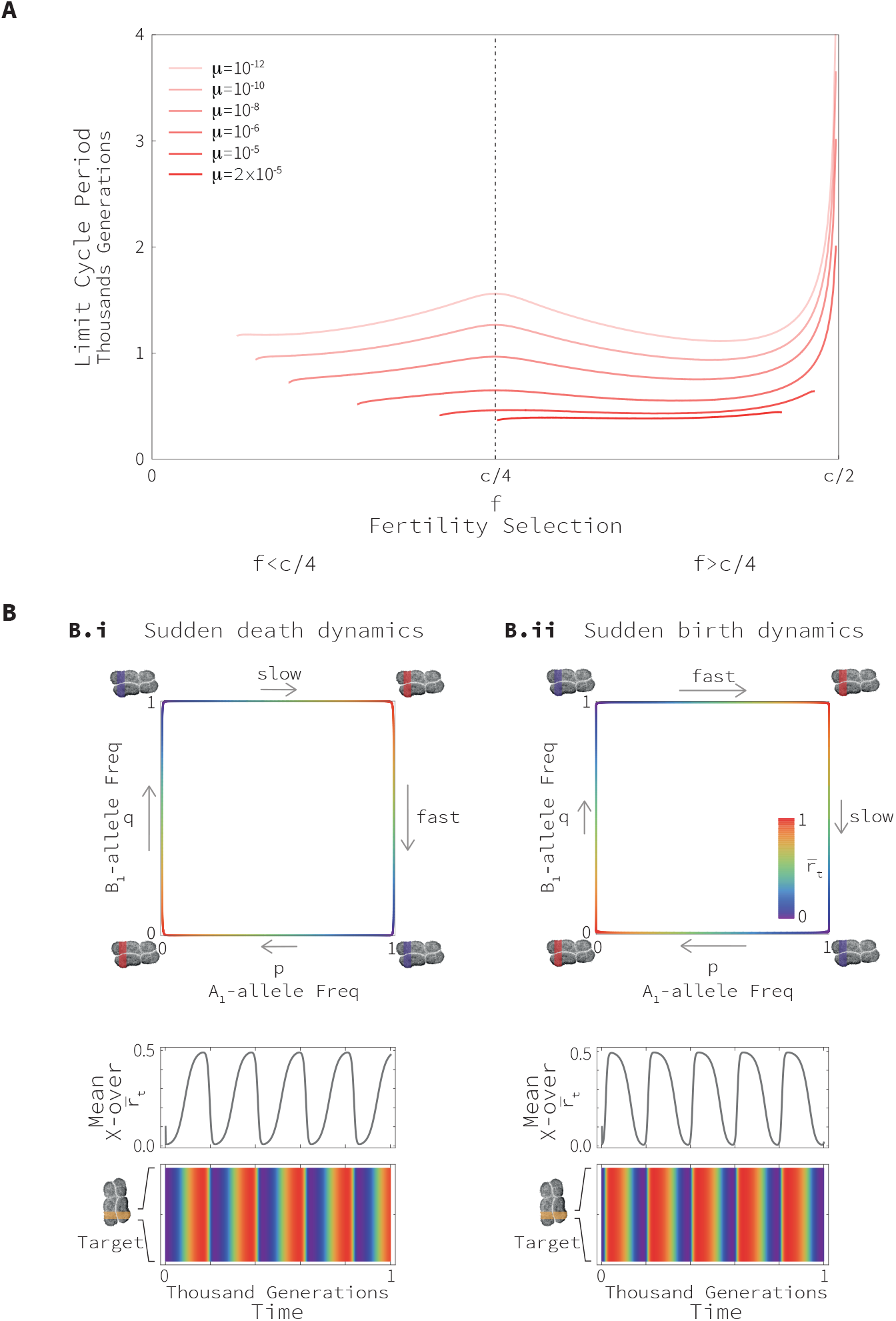
Periodicity of limit cycles. **Panel A** displays the period of limit cycles as a function of the fitness cost *f* within the parameter region 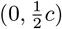. Results are illustrated for different mutation rates *μ* = *μ_A_* = *μ_B_*. The parameter values used to create this figure are *b* = *c* = 1, 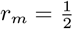. Initial conditions are (*x*_1_(0), *x*_2_(0), *x*_3_(0), *x*_4_(0)) = (*a*, 0, 0,1 – *a*) where *a* = 0.005). **Panel B** displays the genetic and phenotypic oscillations over time. **Sub-panel B.i** displays the case when 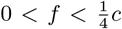. Here the transition from hot to cold is faster than the transition from cold to hot. We refer to this type of dynamics as “sudden death dynamics”. To create this panel we assumed *f* = 0.1. **Sub-panel B.ii** displays the case when 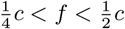. Here the transition from cold to hot is faster than the transition from hot to cold. We refer to this type of dynamics as “sudden birth dynamics”. To create this panel we assumed *f* = 0.4. Within each subpanel there are three figures: The first one depicts the change in allele frequencies in the limit cycle. The second and third figures depict the change in crossover probability at the target locus over time. Heat maps (third figures) are designed so that red colours correspond to high crossover probabilities (hotspot), and blue colours correspond to low crossover probabilities (coldspot). In Panel B we assumed that *b* = *c* = 1, 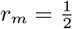, *μ* = 10^-9^, and *p*(0) = 0.99, *q*(0) = 0.01, *D*(0) = 0.

The number of generations required for the replacement of one allele at the target locus (*A*) relative to such replacement at the modifier locus (*B*), depends on the selection strength and the mutation probability. When 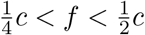, the replacement of alleles at the modifier locus is faster than that at the target locus (Figure 3.B.ii). When 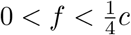, the replacement of alleles at locus *A* is slower than that at locus *B* (Figure 3.B.i). The closer fertility is to 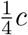, the lower the relative difference between loci in the speed of replacement of alleles. The higher the mutation probability, the lower the relative difference between loci in the speed of replacement of alleles. This difference in the speed of fixation of the alleles in PRDM9 and target loci is relevant because it would allow us to calibrate the model for the strength of fertility selection acting on recombination. In particular, a signature of stronger selective sweep in the PRDM9 locus (relative to target loci) would suggest that the fertility selection *f* is between *c*/4 and *c*/2. On the contrary, a signature of stronger selective sweep in the target loci (relative to the PRDM9 locus) would suggest that the fertility selection *f* is between 0 and *c*/4.

We extend our model to consider a second target locus in order to gain insight into the dynamics when there is more than one target (Section 4 of the SM). In particular, we are interested in exploring whether the oscillation of the mean crossover probability in each target site will be maintained and if it is maintained whether both targets will be oscillating simultaneously (in *synchrony*) or not (in *asynchrony*). We show numerically that starting with populations where one gametic type is almost fixed, the system converges to a limit cycle. In this limit cycle the mean crossover probability at each target loci oscillates asynchronously while the average across target sites (*genome* mean crossover probability) remains close to constant (Figure 4.B). Notice that in the absence of mutation the mean crossover probability at each target locus oscillates in synchrony with each other and thus the genome mean crossover probability changes over time (Figure 4.A). Furthermore, the oscillations stop after a certain amount of time (as in the absence of mutations the dynamics correspond to an heteroclynic cycle).

**Figure 4:**
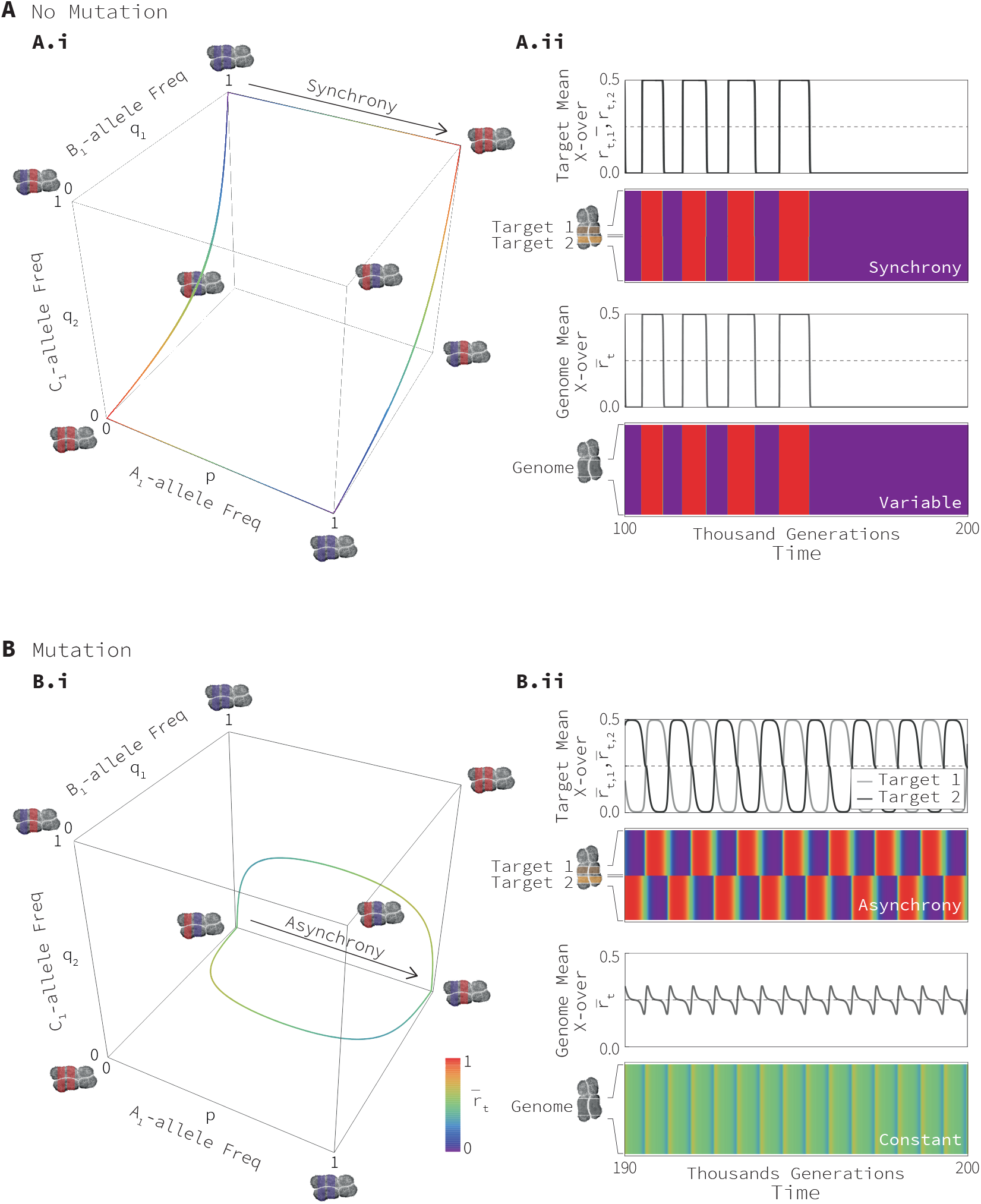
Genetic and phenotypic variation over time in a system with two targets. Cubes represent the change in allele frequency in PRDM9 and target loci. Figures depict the average recombination in each of the target sites and the genome (meaning the average across all targets). These are represented both as lines and heat maps. **Panel A** corresponds to the case where there are no mutations (*μ_A_* = *μ_B_* = *μ_C_* = 0). **Sub-panel A.ii** shows that the dynamics converge to heteroclynic cycles such that both target loci are hot or cold at the same time (synchronic oscillation) and the average genome recombination is variable (oscillating with the average recombination in each target). **Panel B** corresponds to the case where there are mutations (in particular *μ_A_* = 10^-8^, and *μ_B_* = *μ_C_* = 10^-6^). **Sub-panel B.ii** shows that the dynamics converge to limit cycles such that one target is hot when the other is cold or vice-versa (asynchronic oscillation) and the average genome recombination is close to being constant (showing minimal oscillation). Here we assumed that *f* = 0.18, *b* = 1, 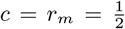, and *p*(0) = *q*_1_(0) = 0.9, *q*_2_(0) = 0.8, *D*(0) = 0.

Interestingly, we find that the mutation probability in PRDM9 and the strength of fertility selection determine the degree of asynchrony between target loci. With mutation in PRDM9 (*μ_A_* > 0), targets oscillate asynchronously with one exhibiting a high mean crossover probability when the other exhibits a low one. Because the mean crossover probability in one target compensates the mean crossover in the other one, the genome mean crossover probability remains close to constant over time. The greater the mutation probability, the greater the asynchrony in mean crossover probability between target sites and the lower the variation in mean crossover probability in the genome (Figure 5). The intuitive reason why is that mutation generates enough genetic variation so that fertility selection can drive the sweep of a hot allele. When the two targets are hot, conversion turns them cold. As they become colder, selection for an alternative PRDM9 allele mounts up until a selective sweep in PRDM9 happens, turning coldspots into hotspots and vice-versa. If rare PRDM9 alleles are at extremely low frequencies, for example when *μ_A_* = 0, selection must mount up considerably —the two targets need to become cold before a selective sweep that turn both of them hot is triggered (Figure 4.A). As a result, targets oscillate synchronously. In contrast, if there is some standing variation in PRDM9 due to mutation (*μ_A_* > 0), such that rare PRDM9 alleles are at low —but not extremely low frequencies—, the selective sweep will be triggered sooner. Because the frequency of hot alleles in the two targets will never be exactly the same, one of them always start to turn cold before the other one does. When the mutation rate is sufficiently high, one cold target is enough to trigger the selective sweep even when the other target is hot (Figure 4.A). As a result, the selective sweep turns the hotspot into a coldspot and the coldspot into a hotspot, with the targets oscillate asynchronously. This logic extends to the effect of fertility selection, *f*, on the asynchrony of the oscillations: as fertility selection increases, the selective sweep can happen without both targets being cold thus favouring asynchronous oscillations.

**Figure 5:**
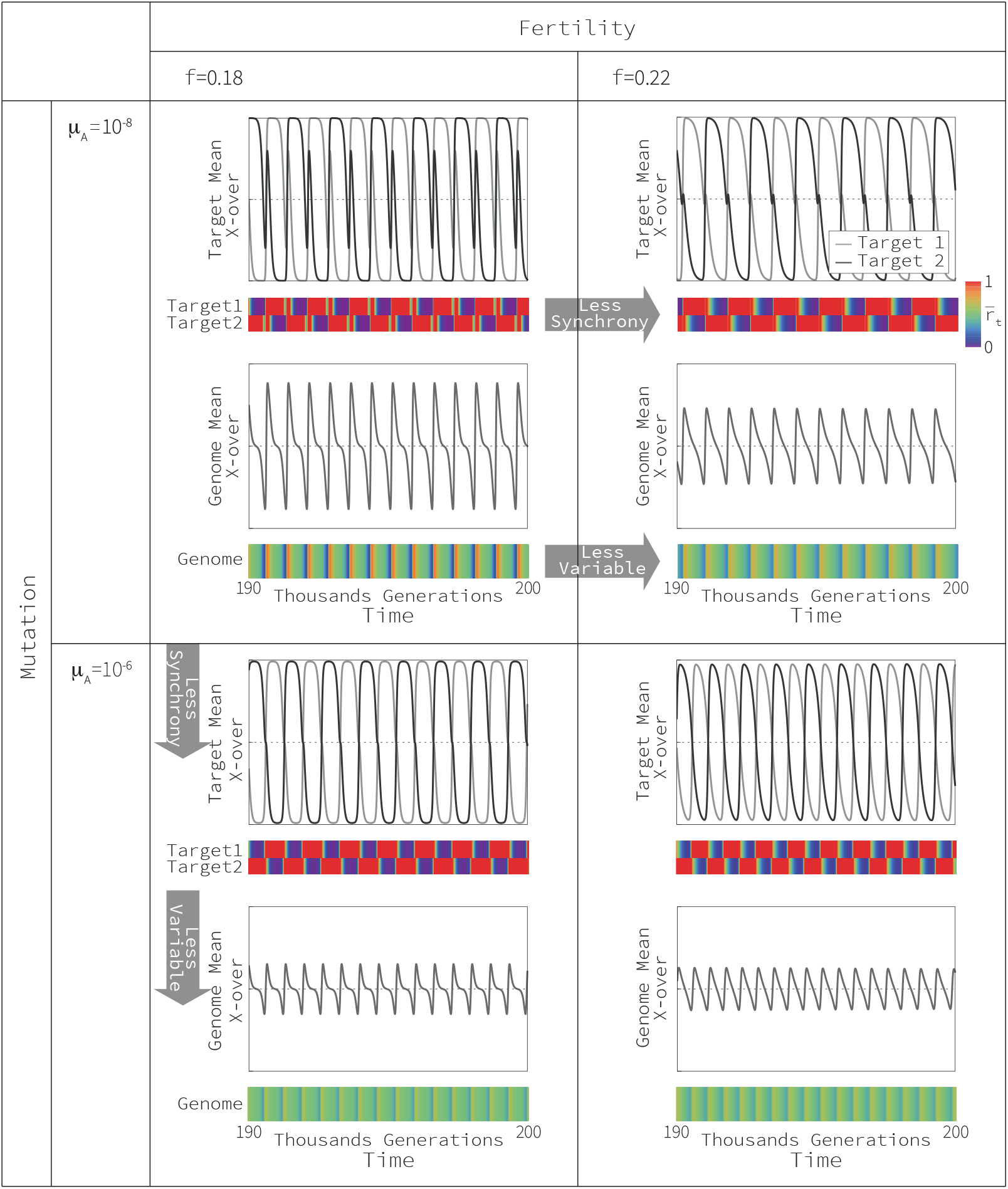
Effect of mutation probability and selection on the asynchrony of oscillations in each target. This figure depicts the change in crossover probability at each of two target loci for different combinations of the mutation probability and fertility selection, along with the genome-wide mean crossover probability. This figure shows that the greater the mutation probability the lower the overlap of hot and cold phenotypes at different target loci (the greater the asynchrony) and the lower the oscillation of the genome mean crossover probability. Notice that mutation does not affect the amplitude of the oscillation in each target. Here we assumed *b* = 1, 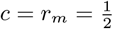, and initial frequencies of: *p*(0) = *q*_1_(0) = 0.9, *q*_2_(0) = 0.8, *D*(0) = 0.

Finally, we formulate a model assuming that the population is finite and that there may be any number of target loci (Section 5 of the SM). We find that the behaviour of the system does not change qualitatively when considering small populations and/or more than two target loci. In particular, we show that the crossover rate in individual target sites oscillates widely while the genome’s crossover rate is more stable (Figure 6). We explore differences in populations of size 5 and 10 thousand individuals (*N* = 5,000 and *N* = 10,000). We find that in both cases the crossover rate in individual target sites oscillates widely, with the turnover of hotspots accelerating as the population size increases (Figure 6). We also explored differences in genomes with two and ten target loci. We find that in both cases the crossover rate in individual target sites oscillates widely, with the turnover of hotspots decelerating as the number of target size increases (Figure 6). The reason why is that, in finite populations, genetic drift favours the loss of rare alleles that need to be recreated again for a selective sweep to occur. As the recreation of alleles is driven by mutations (which are rare), genetic drift lengthens hot and cold phases, increasing the average period of oscillations.

**Figure 6:**
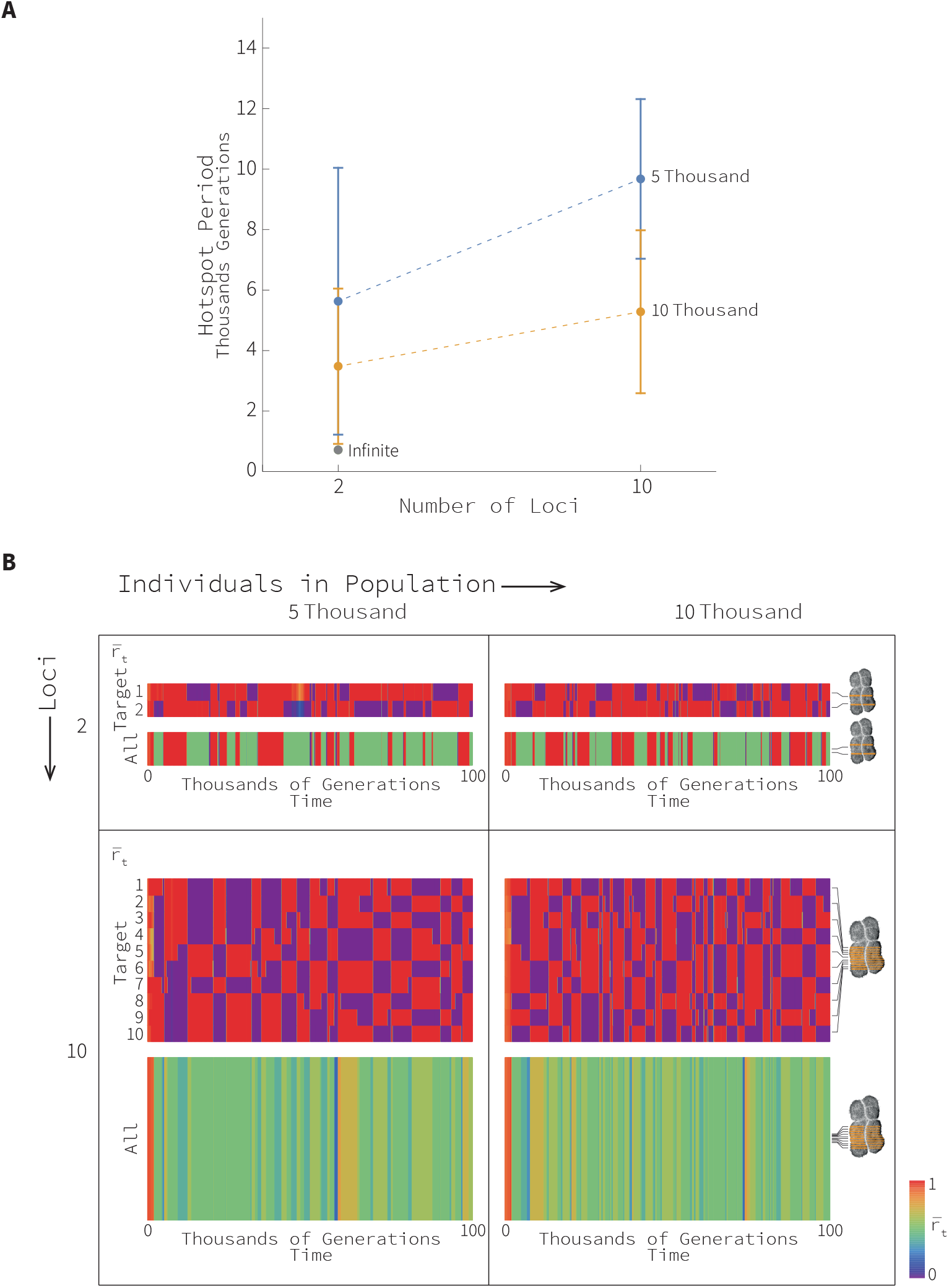
Effect of population sizes and number of targets on the speed of hotspot oscillations in finite populations. **Panel A** shows the average period of an oscillation between hot and cold states. Blue and orange dots represent the average value of the period (for two targets in 100 thousand generations) when the population size is five and ten thousand individuals respectively. The grey dot represents the equivalent value for an infinite population. Vertical bars correspond to the standard deviation. We make the conservative assumption that there is only one attempt to produce one double strand break per genome per generation. **Panel B** shows examples of dynamics. Examples are arranged in a table with five and ten thousand individuals in the columns and two and ten loci in the rows. This figure shows that the smaller the population size or the greater the number of target loci the greater the period of hotspot oscillations. Here we assumed that *f* = 0.4, *b* = *c* = 1, 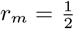 (the recombination rate between any pair of loci is also 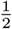), *μ_A_* = 10^-6^, *μ_B_* = 10^-7^, and *p*(0) = 0.95, *q*(0) = 0.95, *D*(0) = 0.

## Discussion

Our model shows that the co-evolutionary dynamics between a gene for target specificity (like PRDM9) and its target are more complex than previously assumed. Our two-locus two-allele model for the interaction of these loci under fertility selection, gene conversion and recurrent mutation shows that there are three qualitatively different outcomes. We find that, when fertility selection is stronger than gene conversion, fertility outcompetes conversion favouring the fixation of target alleles that bind PRDM9 (hot alleles) and recombination hotspots are not eroded by gene conversion —they do not die. We also find that, when fertility selection is weaker than gene conversion, conversion outcompetes fertility favouring target alleles that do not bind PRDM9 (cold alleles) and recombination hotspots are eroded by gene conversion —they die. However, mutations work together with fertility to favour rare PRDM9 alleles that bind targets previously favoured by conversion. In this case there are two possible outcomes: i. when the target site bound by a rare PRDM9 is frequent enough, natural selection favours the re-activation of the former hotspot (resuscitation) which will be followed by its erosion (death) in a never-ending succession of resuscitation-death events —a limit cycle of allelic frequencies resulting in a permanent and regular oscillation between high (hotspot) and low (coldspot) recombination values; ii. when the target allele bound by a rare PRDM9 is not frequent enough, natural selection favours a partial re-activation of the former hotspot followed by a partial erosion until the target settles in a constant intermediate recombination activity —a polymorphic equilibrium of allelic frequencies resulting in a constant recombination value slightly below the average between recombination in hotspot and coldspots (see 1). We find that in our model fertility selection acts in a frequency dependent manner —although it is not explicitly modelled as frequency dependent— with selection for hot alleles being weak when they are abundant but strong when hot alleles are rare.

The co-evolutionary model presented here advances our understanding of the evolutionary dynamics of PRDM9-directed recombination hotspots. Our work advances a previous evolutionary model of PRDM9 alleles (that do not consider the evolutionary feedbacks from their target sites) [21] and a numerical exploration of co-evolutionary models of PRDM9 and their targets [47] by identifying how the interplay between fertility, conversion, and mutation leads to multiple qualitatively different outcomes. Our work also advances a previous co-evolutionary model of PRDM9 and their targets [46] by showing that mutations can lead to permanent and regular oscillations between hotspots and coldspots. Previous work finds that in the absence of mutations hotspots die and resuscitate with the intensity and duration of their hot and cold states ever increasing until, in practical terms, the target becomes stuck in a hot or cold state (see 2 B.i). This result is unrealistic because the formation of recombination hotspots that are not eroded is contrary to empirical observations. Our model shows that in the presence of mutations in PRDM9 and target sites hotspots die and resuscitate with the intensity and duration of their hot and cold states becoming constant (see 2 B.ii). This result is not only more realistic but also allows us to establish how fertility, conversion and mutation can affect the life expectancy of recombination hotspots. Unexpectedly, we find that the life expectancy of hotspots does not always increase with the strength of fertility selection (Figure 3 A). In particular, assuming equal mutation rates in PRDM9 and target sites, the higher the mutation rate and the farther the strength of fertility from 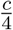, the shorter the life expectancy of recombination hotspots (Figure 3). These predictions can be tested against actual data on the life expectancy of hotspots.

Our model finds that when the strength of fertility selection exceeds one quarter of the conversion probability, 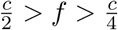, target sites oscillate exhibiting gradual death of hotspots but sudden birth of hotspots (*sudden birth* dynamics) (Figure 3 B.ii). In these cases, our model predicts stronger selective sweeps in genomic regions flanking PRDM9 relative to those flanking recombination hotspots. When the strength of fertility selection falls below one quarter of the conversion probability, 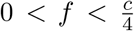, recombination sites oscillate with sudden death of hotspots but gradual birth of hotspots (*sudden death* dynamics) (Figure 3 B.i). In these cases, our model predicts weaker selective sweeps in genomic regions flanking PRDM9 relative to those flanking recombination hotspots. Our work suggests the possibility of using the genetic variability in genomic regions flanking PRDM9 relative to that near recombination hotspots to calibrate the strength of fertility selection acting on human populations.

Like any other model, the model we present here makes some simplifying assumptions that do not correspond to the reality of what is being modelled. In order to understand the interplay of the three evolutionary forces we are interested in —fertility selection, gene conversion and recurrent mutation— we assume that PRDM9 acts on a single target and both PRDM9 and target sites can carry two alleles only. In reality, both PRDM9 and target loci can be segregating multiple alleles. Furthermore, each PRDM9 allele can be acting on hundreds of targets that carry the same sequence motif (family of target sites). However, understanding the role of each of the evolutionary forces considered allows us to gain some insight on the extension to multiple alleles in each loci and multiple targets. Namely, when fertility selection is weaker than conversion, our model finds that fertility selection drives to fixation a PRDM9 binding a target (hot allele) when the motif recognised is close to fixation. In this case selection results in the formation of a recombination hotspots. Once fertility drives the formation of this recombination hotspot, conversion erodes the motif turning the target into a recombination coldspot again. However, our model also finds that fertility and conversion balance the frequencies of PRDM9 binding a target (hot allele) and PRDM9 not binding that target (cold allele) when the motif recognised is not close to fixation. In this case selection results in the formation of a target with intermediate recombination activity. This target exhibiting mild recombination does not suffer erosion from conversion and its recombination activity remains constant over time. Our model shows that how close to fixation the new binding motif should be is a function of the relative strength of fertility selection and the mutation rate (see 1). These conclusions can be extended to multiple allele and targets. If instead of one target each allele is acting on multiple targets, our model suggests that fertility selection will favour the resuscitation of multiple recombination hotspots when the motif binding the PRDM9 is close to fixation. If instead of two alleles PRDM9 is segregating multiple alleles each acting on a different family of targets, our model suggests that fertility selection will favour the birth of multiple recombination hotspots when the motif biding the PRDM9 is close to fixation. This will be true as long as the new alleles in PRDM9 are acting on new targets that do not cause a fitness host in the carrier by causing a break in a functional gene, for example. However, when having multiple targets the fertility cost caused by a loss of one recombination hotspot should be smaller than in the one target model we explored here. Therefore we expect that fertility will drive the resuscitation or birth of recombination hotspots only when sufficient number of recombination hotspots have been turned into coldspots.

An exhaustive study of the dynamics of all possible multi-allelelic multi-target extensions of our model is beyond the scope of this work. However, to test the validity of our insights, we did extend our deterministic two-locus one target model model to explore the impact of a second target. When fertility selection is weaker than conversion and initially one gametic type is near fixation, permanent and regular oscillations between hotspots and coldspots are observed in both target sites. In principle, the two target sites can be hot and cold at the same time (oscillate synchronously) or one target site can be hot when the other is cold (oscillate asynchronously). When both targets oscillate synchronously the average recombination across targets (the genome’s recombination) oscillates with them. When both targets oscillate asynchronously the genome’s recombination remains constant. Interestingly, we find that mutations are key in maintaining a constant recombination rate across the genome while permitting the recombination rate in each site to oscillate widely (Figure). We find that the greater the mutation in PRDM9 the smaller the fluctuation of recombination across the genome (Figure 5). The greater the fertility selection the smaller the fluctuation of recombination across the genome (Figure 5). The intuitive reason is that the recombination rate in one site conditions the strength of selection in the other sites. The more recombination coldspots, the greater the strength of fertility selection necessary to resuscitate other hotspots. The greater the availability of matching PRDM9 mutants the smaller the strength of fertility selection necessary to resuscitate other hotspots. Therefore, with lower mutation probability and/or fertility cost all target sites need to be cold before selection favours a switch resulting in all of them being hot (synchronous oscillation). With higher mutation probability and/or fertility cost some target sites need to be cold before selection favours a switch resulting in some of them being hot while the other becomes cold (asynchronous oscillation). In the absence of mutation all target loci oscillate synchronously and the recombination rate in the genome oscillates widely.

To further explore the validity of our insights we formulated a finite population (stochastic) version of our model with multiple alleles and multiple target sites. We explore populations of a size similar to the expected population size in humans, that is *N* = 5, 000 and N = 10,000 individuals. We consider one PRDM9 and multiple targets. In first instance we consider the case whenPRDM9 segregates 2 alleles and all targets segregate the same pair of motifs, that is all targets are part of the same family. We find that when fertility is lower than conversion and initially multiple motifs are rare, the crossover rate in individual target oscillates widely while the genome’s crossover is more stable (Figure 6). In addition we consider the case when PRDM9 segregates 4 alleles and targets equally split into two families that segregate two different alleles each. Initially, PRDM9 segregates only two alleles matching the alleles segregating at one first family. We observe that the crossover rate in individual targets of this family oscillate widely. When the two alleles matching the motifs of the second family are introduced. We observe that the crossover rate in individual targets of this family also oscillate widely and PRDM9 alternates between the use of the two families (Figure 8). This is true as long as both families do not incur in any fitness costs like, for example, having one of its target motifs present in a functional gene and thus inducing a break in this gene. When one family has a greater probability of incurring in a fitness cost relative to the other family, the PRDM9 alleles that target the former family will be purged from the population (Figure 8).

**Figure 7:**
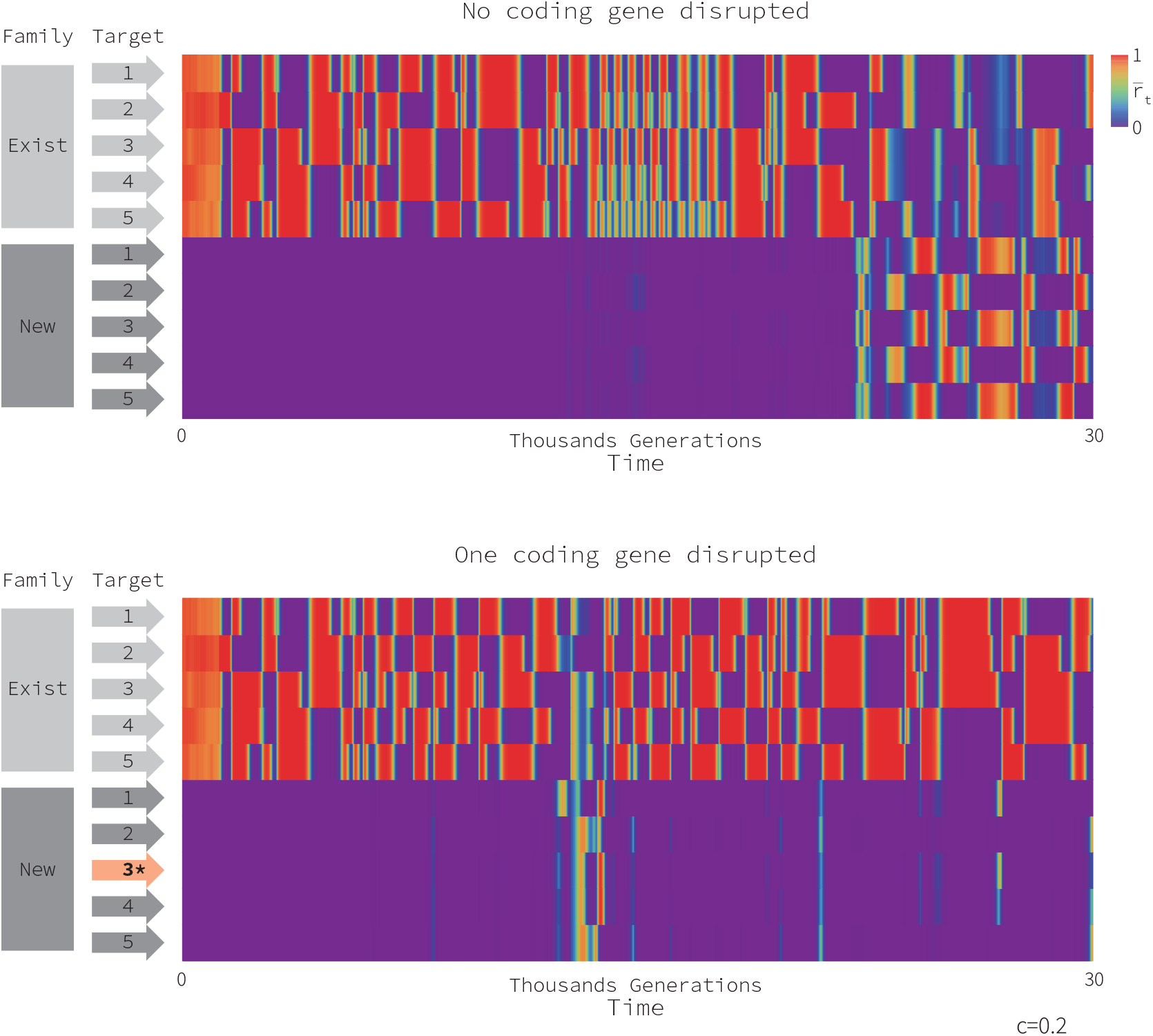
Recombination Hotspots dynamics with crossing-over costs. We consider here a model with 10 target loci separated into two families. First five target loci bear sequences B1 and B2; crossing-over in these locations is never detrimental. The other five target loci bear sequences B3 and B4; crossing-over on target locus 8 bears no cost in **Panel A** and a cost 0.2 (g = 0.2) in **Panel B**. The first 1,000 generations are a burn-out phase were everything is neutral. Until generation 5,000, PRDM9 locus can only target sequences B1 and B2; target loci 6 to 10 can never be targeted. After generation 5,000, mutations on PRDM9 can create alternative alleles able to target all B alleles, that is, all target loci. Parameter values include: 10,000 individuals, *b* = *c* = 1, *f* = 0.4, *μ_A_* = 10^-^05 and *μ_B_* = 10 – 6. All loci are assumed to be freely recombining. Population is initialised with *p* = 0.99, *q* = 0.95 for all loci of the first family of target loci, and *q* = 0.5 (as we assume they are initially neutral) for all loci of the second family of target loci. Note that the second family of target loci evolve neutrally during the first 5,000 generations.

**Figure 8:**
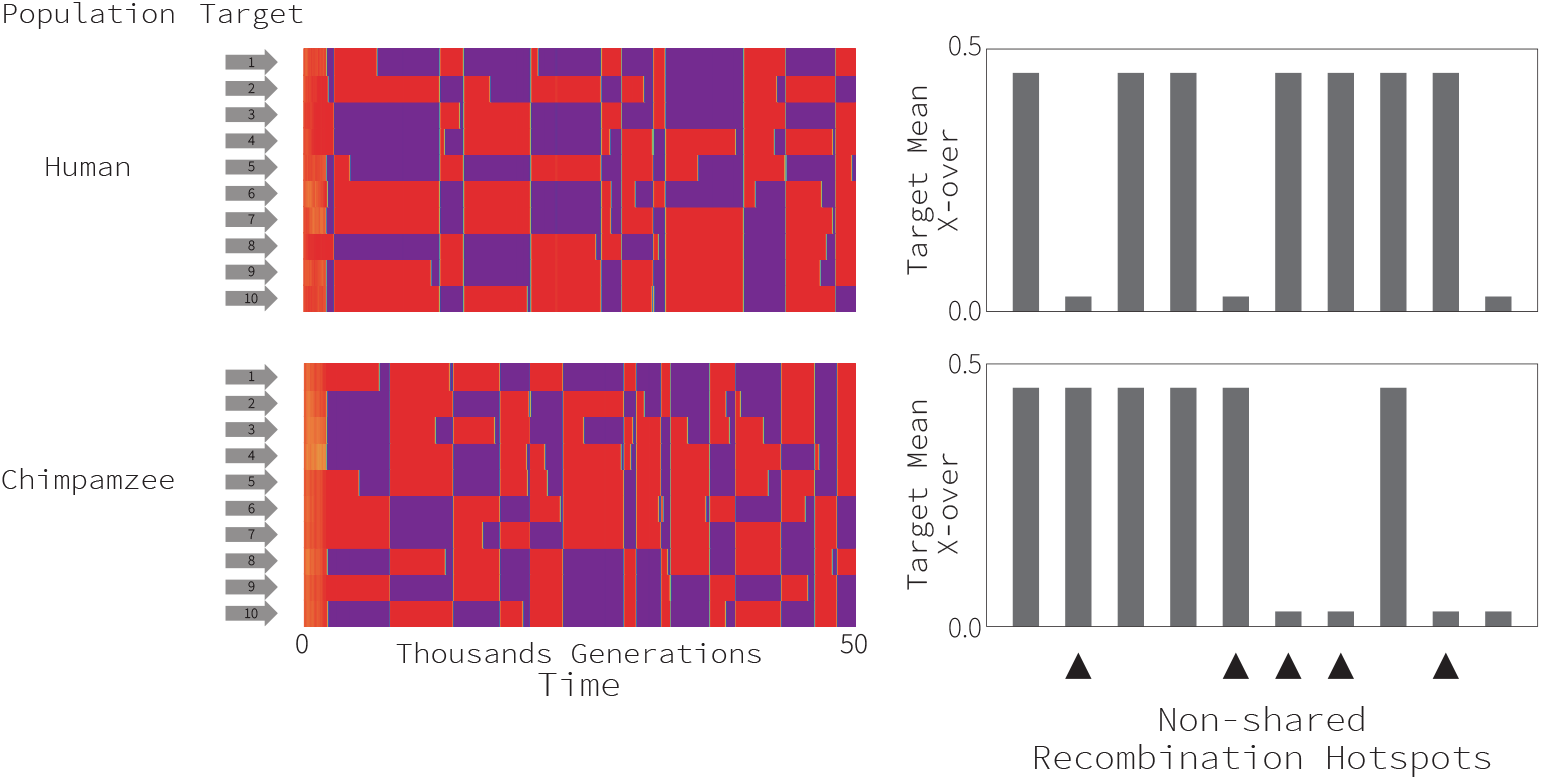
Evolution of the recombination landscape in two separate populations when hotspots resuscitate. This figure shows the hotspots that are shared and non-shared between these two populations after 10 thousand generations. Here we assume that *b* = *c* = 1, 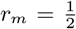 (the recombination rate between any pair of loci is also 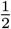), *μ_A_* = 10^-6^, *μ_B_* = 10^-7^, and *p*(0) = 0.95, *q*(0) = 0.95, *D*(0) = 0.

Our work contributes to solving the Recombination Hotspot Paradox by reconciling the self-destructive nature of PRDM9-directed recombination hotspots with the formation of hotspots to maintain the elevated number of such hotspots observed in various genomes [6]. The findings of our model are consistent with all salient features of recombination hotspots namely the death of individual hotspots, the preservation of hotspots at the genome level, and the rapid change of the recombinational landscape (with hotspots rarely being shared between closely related species). We find that when fertility is weaker than conversion, recombination hotspots are eroded by conversion and individual hotspots die. Given the same selection regime when recombination activity is low, fertility favours the formation of recombination hotspots. The formation of hotspot will take place in either families of targets that used to be hotspots, or new families of targets with limited variability of motifs that are not detrimental to the fitness of their carrier. Eventually, a finite number of segregating alleles at PRDM9 should determine an alternation between a finite set of families of target sites families that are cost free for their carrier. Notice that while in our model hotspots are recycled, the location of active hotspots in a particular moment in time should be unique to each subpopulation even within a family of targets (Figure 7). Besides offering a plausible solution to the paradox, our research offers the possibility of calibrating the model we present against genomic data. Such calibration would allow us to provide quantitative predictions of the life expectancy of hotspots given different effects of recombination on fertility. This, in turn, can be used to study the signature of oscillatory recombination hotspots on linkage disequilibrium in the mammalian genome, a pattern that is relevant to better understand links between alleles and diseases.

## Acknowledgements

[FU] thanks Deborah Charlesworth and Brian Charlesworth for details comments and guidance [FU] and [FF] are supported by a NSFDEB-NERC Research Grant NE/T009322/1.

## Author Contributions

All authors contributed equally

