## Supplemental Material for "Co-evolution between PRDM9 and target sites and the Recombination Hotspot Paradox"

### 1. Deterministic one-target model: gamete frequency dynamics

#### *1.1. Conversion and selection*

##### *1.1.1. Two-locus multi-allelic model*

We use a classical deterministic model to describe the change of gamete frequencies under selection, recombination, and mutation in a sufficiently large population such that random genetic drift can be ignored. We assume a randomly mating population of diploid individuals with discrete, non-overlapping generations.

We model the interaction between two loci. The first locus corresponds to a gene (PRDM9-like) coding for a protein that may recognise a specific motif at a second target locus where recombination may be initiated. This modifier locus $A$ (PRDM9-like gene) may carry alleles $A_1, A_2, ..., A_I$, each encoding a protein that attempts to bind a motif at a target locus $B$. Locus $B$ may carry alleles $B_1, B_2, ..., B_K$, each corresponding to a base pair motif that the protein produced by locus $A$ may attempt to bind. In each generation, both modifier alleles in each diploid individual show the same level of expression. One protein from this pool is chosen at random (each type of protein having equal probability $\frac{1}{2}$ of being chosen) and attempts to bind one of the two target motifs at random (each target motif having equal probability $\frac{1}{2}$ of experiencing a binding attempt). We assume that there is only one binding attempt. Therefore, in an individual with genotype $\frac{A_i B_k}{A_j B_l}$,

*Corresponding author

four potential binding attempts can occur ($A_i \rightarrow B_k$, $A_i \rightarrow B_l$, $A_j \rightarrow B_k$, $A_j \rightarrow B_l$), each with equal probability $\frac{1}{4}$. The binding attempt of protein $A_i$ to motif $B_k$ results in a successful binding and a double-strand break of allele $B_k$ with probability $b_{i,k}$. The binding attempt results in a failed binding and a lack of a double-strand break with probability $1 - b_{i,k}$ (where $0 < b_{i,k} < 1$) (Figure S1).

A double-strand break (DSB) initiates recombination and the chromatid that breaks is repaired using its homologous chromatid as a template (1, 2). During the repair process there might be a crossover event in or near the target locus with probability $r_t$ (henceforth crossover probability) and none with probability $1 - r_t$ (where $0 < r_t < 1$) (1, 2). In our model, we assume that modifier and target loci are either far apart in the same chromosome or in separate chromosomes. From a modelling perspective, this means that modifier and target loci always experience recombination independently of whether there is a DSB at the target locus or not, that is the recombination probability is $r_m = \frac{1}{2}$. This assumption is consistent with empirical observations and previous work on the topic (3, 4). During the repair process the allelic motif that breaks is converted into the allelic motif that does not break with probability $c$ and is restored to the allelic motif that breaks with probability $1 - c$ (where $0 < c < 1$) (5, 6, 1, 2). Notice that biased gene conversion results in the over-transmission of the allele that is less likely to break (7, 2) (Figure S1).

Recombination ends up with Mendelian segregation of alleles into gametes. Following previous models (3, 4), we assume that individuals undergoing recombination at the target locus have proper chromosomal segregation and do not suffer any fitness cost, while individuals that do not undergo recombination at the target locus may have defective chromosomal segregation producing aneuploid (non-viable) gametes with probability $f$ (where $0 < f < 1$). Therefore, the fitness of individuals experiencing a recombination event at the target locus is 1 but the fitness of individuals not experiencing a recombination event is $1 - f$ (Figure S1).

Let $x_{i,k}$ be the frequency of type $A_iB_k$ in gametes. Notice that $0 \leq x_{i,k} \leq 1$ and $\sum_{i,k} x_{i,k} = 1$. Random union of gametes results in an embryo with ordered genotype $\frac{A_iB_k}{A_jB_l}$ with frequency $x_{i,k}x_{j,l}$. The probability that this embryo reaches adulthood is independent of its genotype, but its genotype determines the outcome of meiosis in adults. The probability that during meiosis the protein produced by the modifier locus breaks target $B_k$ is $\bar{b}_{ij,k} = \frac{1}{2}(b_{i,k} + b_{j,k})$ because only one of the proteins produced by the two modifier alleles attempts a break. Analogously, the probability that the modifier locus breaks target $B_l$ is $\bar{b}_{ij,l} = \frac{1}{2}(b_{i,l} + b_{j,l})$. Therefore, the probability that the protein produced by the modifier locus breaks one of the two targets (either $B_k$ or $B_l$ is $\bar{\bar{b}}_{ij,kl} = \frac{1}{2}(\bar{b}_{ij,k} + \bar{b}_{ij,l})$ because only one of the two sequence motifs at the target locus can break. Notice that all along we assume that at most one double-strand break can occur. The probability that during meiosis a double-strand break is followed by a crossover event between the flanking regions at the target locus $B$ is $r_t$, and the probability that the motif that breaks is converted into the motif that does not break is $c$. Double-strand break at the target locus is followed by correct Mendelian segregation of types into gametes but, in the absence of DSBs, segregation of gametic types is incorrect with probability $f$. Finally, the probability that during meiosis alleles at the $A$ and $B$ locus experience recombination is $r_m$, which is assumed to be equal to $\frac{1}{2}$ in our numerical analysis. Gametic type segregation brings us back to the beginning of the census (Figure S1).

The frequency of type $A_iB_k$ in gametes after recombination and selection is

$$\begin{aligned} x_{i,k}^{(rs)} &= \frac{1}{\bar{w}} \sum_{j,l} \Big[ \Big( \bar{\bar{b}}_{ij,kl} + (1 - \bar{\bar{b}}_{ij,kl})\,(1-f) \Big)\, x_{i,k}\, x_{j,l} \\ &\quad -c\,\frac{1}{4}\left(\bar{b}_{ij,k}\, x_{i,k}\, x_{j,l} - \bar{b}_{ij,l}\, x_{i,l}\, x_{j,k}\right) \\ &\quad -r_m\, \Big( (1-c)\,\bar{\bar{b}}_{ij,kl} + (1-\bar{\bar{b}}_{ij,kl})\,(1-f) \Big)\,(x_{i,k}\, x_{j,l} - x_{i,l}\, x_{j,k}) \Big] \,, \end{aligned} \tag{1}$$

where

$$\bar{w} = \sum_{i,k} \sum_{j,l} \left[ \bar{\bar{b}}_{ij,kl} + (1 - \bar{\bar{b}}_{ij,kl})(1 - f) \right] x_{i,k}\, x_{j,l} \tag{2}$$

is the population mean fitness.

These changes in gametic type frequency underpin changes in the population mean crossover probability at the target locus,

$$\bar{r}_t = \frac{1}{2} \sum_{i,k} \sum_{j,l} \bar{\bar{b}}_{ij,kl}\, r_t\, x_{i,k}\, x_{j,l}\,, \tag{3}$$

which is the phenotype whose evolution we are interested in.

#### *1.1.2. Two-locus two-allele model*

We consider the above model in the particular case when there are two alleles $(A_1, A_2)$ at the modifier locus and two alleles $(B_1, B_2)$ at the only target locus, resulting in four different gametic types $(A_1B_1, A_1B_2, A_2B_1, A_2B_2)$. Henceforth, we assume that a match between the subscripts of the modifier allele producing the protein that attempts to bind and the allelic sequence that is the target of this attempt to bind results in a double-strand break with probability $b$ (where $0 < b < 1$) and a mismatch between the subscripts prevents a double-strand break. For our modelling purposes this translates into

$$b_{i,k} = \begin{cases} b & \text{if } i = k \\ 0 & \text{if } i \neq k. \end{cases}$$

Notice that two of these gametic types $(A_1B_1, A_2B_2)$ correspond to types producing a protein that matches its own target sequence (recombination enabling types) and the other two $(A_1B_2, A_2B_1)$ correspond to gametic types producing a protein that does not match its own target sequence (recombination disabling types).

The dynamic system describing the change in frequency over time of each of these haplotypes can be obtained from replacing generic subscripts $i$ and $k$ by specific subscripts 1 and 2 in equation (1). The frequency of the gametic type $A_iB_k$ after recombination and selection is

$$
\begin{aligned}
\bar{w}\, x_{1,1}^{(rs)} &= \left(\tfrac{1}{2}\, b + (1 - \tfrac{1}{2}\, b)\,(1 - f) + \tfrac{1}{2}\, b\, f\, x_{1,1} - \tfrac{1}{4}\, b\, c\, x_{1,2}\right)\, x_{1,1} \\
&\quad - \left(\tfrac{1}{2}\, b\, \left(\tfrac{1}{4}\, c + (1 - c)\, r_m\right) + r_m\,(1 - \tfrac{1}{2}\, b)\,(1 - f)\right)\, D\,, \\
\bar{w}\, x_{1,2}^{(rs)} &= \left(\tfrac{1}{2}\, b + (1 - \tfrac{1}{2}\, b)\,(1 - f) - \tfrac{1}{2}\, b\, f\, x_{1,2} + \tfrac{1}{4}\, b\, c\, x_{1,1}\right)\, x_{1,2} \\
&\quad + \left(\tfrac{1}{2}\, b\, \left(\tfrac{1}{4}\, c + (1 - c)\, r_m\right) + r_m\,(1 - \tfrac{1}{2}\, b)\,(1 - f)\right)\, D\,, \\
\bar{w}\, x_{2,1}^{(rs)} &= \left(\tfrac{1}{2}\, b + (1 - \tfrac{1}{2}\, b)\,(1 - f) - \tfrac{1}{2}\, b\, f\, x_{2,1} + \tfrac{1}{4}\, b\, c\, x_{2,2}\right)\, x_{2,1} \\
&\quad + \left(\tfrac{1}{2}\, b\, \left(\tfrac{1}{4}\, c + (1 - c)\, r_m\right) + r_m\,(1 - \tfrac{1}{2}\, b)\,(1 - f)\right)\, D\,, \\
\bar{w}\, x_{2,2}^{(rs)} &= \left(\tfrac{1}{2}\, b + (1 - \tfrac{1}{2}\, b)\,(1 - f) + \tfrac{1}{2}\, b\, f\, x_{2,2} - \tfrac{1}{4}\, b\, c\, x_{2,1}\right)\, x_{2,2} \\
&\quad - \left(\tfrac{1}{2}\, b\, \left(\tfrac{1}{4}\, c + (1 - c)\, r_m\right) + r_m\,(1 - \tfrac{1}{2}\, b)\,(1 - f)\right)\, D\,,
\end{aligned}
\tag{4}
$$

where

$$
\bar{w} = \tfrac{1}{2}\, b + (1 - \tfrac{1}{2}\, b)\,(1 - f) + \tfrac{1}{2}\, b\, f\,(x_{1,1}^2 + x_{2,2}^2 - x_{1,2}^2 - x_{2,1}^2) \tag{5}
$$

is the population mean fitness, and

$$
D = x_{1,1}\, x_{2,2} - x_{1,2}\, x_{2,1} \tag{6}
$$

is the linkage disequilibrium.

To simplify we define parameters $\alpha, \beta, \gamma$, and $\delta$ as follows:

$$\begin{aligned}
\alpha &= \tfrac{1}{2}\, b + \left(1 - \tfrac{1}{2}\, b\right) (1 - f) \ , \\
\beta &= \tfrac{1}{2}\, b\, f \, , \\
\gamma &= \tfrac{1}{4}\, b\, c \, , \\
\delta &= \tfrac{1}{2}\, b \, \left(\tfrac{1}{4}\, c + (1 - c)\ r_m\right) + r_m \left(1 - \tfrac{1}{2}\, b\right)\ (1 - f) \ ,
\end{aligned} \tag{7}$$

which allows us to re-write the system of equations (4) as follows:

$$\begin{aligned}
\bar{w}\, x_{1,1}^{(rs)} &= (\alpha + \beta\, x_{1,1} - \gamma\, x_{1,2})\, x_{1,1} - \delta D \, , \\
\bar{w}\, x_{1,2}^{(rs)} &= (\alpha - \beta\, x_{1,2} + \gamma\, x_{1,1})\, x_{1,2} + \delta D \, , \\
\bar{w}\, x_{2,1}^{(rs)} &= (\alpha - \beta\, x_{2,1} + \gamma\, x_{2,2})\, x_{2,1} + \delta D \, , \\
\bar{w}\, x_{2,2}^{(rs)} &= (\alpha + \beta\, x_{2,2} - \gamma\, x_{2,1})\, x_{2,2} - \delta D \, ,
\end{aligned} \tag{8}$$

with population mean fitness

$$\bar{w} = \alpha + \beta(x_{1,1}^2 - x_{1,2}^2 - x_{2,1}^2 + x_{2,2}^2) \, . \tag{9}$$

Note that $0 < \alpha, \beta, \gamma, \delta < 1$ and

$$\alpha > \beta, \ \alpha > \gamma, \ \text{and} \ \tfrac{1}{2}\gamma \leq \delta \, . \tag{10}$$

Let subscripts 1, 2, 3, and 4 denote each of the four elements of a vector $\mathbf{x}$. Notice that using this notation, subscripts 1,2,3, and 4 refer to haplotypes $(1,1)$, $(1,2)$, $(2,1)$, and $(2,2)$ respectively. Thus, $\mathbf{x} = (x_1, x_2, x_3, x_4)^\top$ is the (column) vector of state variables ($^\top$ denotes transposition). The frequencies of the gametic types after recombination and selection are given by

$$\bar{w}\,\mathbf{x}^{(rs)} = (\mathbf{W}\mathbf{x}) \circ \mathbf{x} - \mathbf{d}\,\delta D\,, \tag{11}$$

where

$$\mathbf{W} = \begin{pmatrix} \alpha+\beta & \alpha-\gamma & \alpha & \alpha \\ \alpha+\gamma & \alpha-\beta & \alpha & \alpha \\ \alpha & \alpha & \alpha-\beta & \alpha+\gamma \\ \alpha & \alpha & \alpha-\gamma & \alpha+\beta \end{pmatrix} \tag{12}$$

is the fitness matrix, $\mathbf{d} = (\,1, -1, -1,\,1)^{\top}$, and $\circ$ denotes the pointwise product of vectors or matrices.

Using the original variables, we can write this matrix as

$$\mathbf{W} = \begin{pmatrix} 1-f(1-b) & 1-f(1-\frac{b}{2})-\frac{bc}{4} & 1-f(1-\frac{b}{2}) & 1-f(1-\frac{b}{2}) \\ 1-f(1-\frac{b}{2})+\frac{bc}{4} & 1-f & 1-f(1-\frac{b}{2}) & 1-f(1-\frac{b}{2}) \\ 1-f(1-\frac{b}{2}) & 1-f(1-\frac{b}{2}) & 1-f & 1-f(1-\frac{b}{2})+\frac{bc}{4} \\ 1-f(1-\frac{b}{2}) & 1-f(1-\frac{b}{2}) & 1-f(1-\frac{b}{2})-\frac{bc}{4} & 1-f(1-b) \end{pmatrix}. \tag{13}$$

We note that this matrix has the following structure:

$$\mathbf{W} = \big(\mathbf{U} - \big(\mathbf{U} - \mathbf{B}\big) \circ \mathbf{U}f + \mathbf{B} \circ \mathbf{C}\big)\,, \tag{14}$$

where $\mathbf{U}$ is a four by four matrix of ones,

$$\mathbf{B} = \tfrac{1}{2}b \begin{pmatrix} 2 & 1 & 1 & 1 \\ 1 & 0 & 1 & 1 \\ 1 & 1 & 0 & 1 \\ 1 & 1 & 1 & 2 \end{pmatrix} \tag{15}$$

is the break matrix, and

$$\mathbf{C} = \tfrac{1}{2}c \begin{pmatrix} 0 & -1 & 0 & 0 \\ 1 & 0 & 0 & 0 \\ 0 & 0 & 0 & 1 \\ 0 & 0 & -1 & 0 \end{pmatrix} \tag{16}$$

is the conversion matrix. Here, $\circ$ denotes the Schur (pointwise) product of matrices.

#### *1.2. Conversion, selection and mutation*

##### *1.2.1. Two-locus multi-allelic model*

We now introduce mutations in our model in both modifier and target loci. We denote the mutation probability, from $A_i \to A_j$ by $\mu_{A,ij}$, where $i \neq j$. Then $\mu_{A,ii} = 1 - \sum_{j:j\neq i} \mu_{A,ij}$ is the probability that $A_i$ does not mutate. Analogously, we denote the mutation probabilities at locus $B$ by $\mu_{B,kl}$. Of course, we assume that the mutation probabilities are sufficiently small, so that the probabilities that no mutation occurs are close to 1 for all alleles $A_i$ and $B_k$. Then the probability that the gametic type $A_jB_l$ changes to $A_iB_k$ as a result of a mutation is $\mu_{A,ji}\,\mu_{B,lk}$. Therefore, the frequencies of gametes after recombination, selection, mutation and reproduction, i.e., at the beginning of the next generation, are

$$x'_{i,k} = \sum_{j,l} x^{(rs)}_{j,l} \mu_{A,ji}\,\mu_{B,lk}\,, \tag{17}$$

where $x^{(rs)}_{j,l}$ is given in eq. (1).

If mutation is very weak so that terms of higher order than 1 in $\mu_{A,ji}$ and $\mu_{B,lk}$ can be ignored, then

$$\mu_{A,ji}\,\mu_{B,lk} \approx 0 \qquad \text{if } i \neq j \text{ and } l \neq k\,, \tag{18a}$$

$$\mu_{A,ii}\,\mu_{B,lk} \approx \mu_{B,lk} \qquad \text{if } l \neq k\,, \tag{18b}$$

$$\mu_{A,ij}\,\mu_{B,kk} \approx \mu_{A,ij} \qquad \text{if } i \neq j\,, \tag{18c}$$

$$\mu_{A,ii}\,\mu_{B,kk} \approx 1 - \sum_{j:j\neq i} \mu_{A,ij} - \sum_{l:l\neq k} \mu_{B,kl}\,. \tag{18d}$$

This yields

$$x'_{i,k} \approx x^{(rs)}_{i,k}\Big(1 - \sum_{j:j\neq i} \mu_{A,ij} - \sum_{l:l\neq k} \mu_{B,kl}\Big) + \sum_{j:j\neq i} x^{(rs)}_{j,k}\mu_{A,ji} + \sum_{l:l\neq k} x^{(rs)}_{i,l}\mu_{B,lk}\,. \tag{19}$$

The mutation matrix of the full system is the Kronecker product of the mutation matrices $(\mu_{A,ij})$ and $(\mu_{B,kl})$ at the individual loci. An important special case is the one in which all mutation probabilities at a locus are identical.

#### *1.2.2. Two-locus two-allele model*

We now introduce mutations in the 2-allele version of the model. For simplicity we assume that the mutation probability from allele $A_1$ to $A_2$ is the same as from $A_2$ to $A_1$ and designate it $\mu_A$. Analogously, we assume that the mutation probability from $B_1$ to $B_2$ is the same as from $B_2$ to $B_1$ and denote it by $\mu_B$. From eqs. (11) and (17) we infer that the recursion equations for the gamete frequencies are given by

$$\bar{w}\,\mathbf{x} = \mathbf{M}\big((\mathbf{W}\mathbf{x}) \circ \mathbf{x} - \mathbf{d}\,\delta D\big)\,, \tag{20}$$

where $\mathbf{M}$ is the mutation matrix resulting from the Kronecker product of the mutation matrices at loci $A$ and $B$,

$$\mathbf{M}_A = \begin{pmatrix} 1-\mu_A & \mu_A \\ \mu_A & 1-\mu_A \end{pmatrix} \text{ and } \mathbf{M}_B = \begin{pmatrix} 1-\mu_B & \mu_B \\ \mu_B & 1-\mu_B \end{pmatrix}, \tag{21}$$

i.e., $\mathbf{M} = \mathbf{M}_A \otimes \mathbf{M}_B$. The population mean fitness $\bar{w}$ is unaffected by mutation.

From eqs. (8) and (20), we immediately infer that the dynamical equations remain unchanged if we divide these equations by $\alpha$. Therefore, without loss of generality, we can set $\alpha = 1$ by applying the rescaling $\beta \to \beta/\alpha$, $\gamma \to \beta/\alpha$, and $\delta \to \delta/\alpha$ and setting

$$\bar{w} = 1 + \beta(x_{1,1}^2 - x_{1,2}^2 - x_{2,1}^2 + x_{2,2}^2) \tag{22}$$

(with the new $\beta$). For the rest of this paper, we assume $\alpha = 1$ and eq. (22).

### 2. Deterministic one target model: allele frequency and linkage disequilibrium dynamics

It will often be more convenient, and yield more insight, to perform the analysis of the two-locus two-alleles model in terms of the allele frequencies, $p = x_1 + x_2$ of $A_1$ and $q = x_1 + x_3$ of $B_1$, and the linkage disequilibrium, $D$, instead of the gamete frequencies.

#### *2.1. Conversion and selection*

The following is the transformation from $(p, q, D)$ coordinates to $(x_1, x_2, x_3, x_4)$ coordinates

$$\begin{aligned} x_1 &= pq + D \\ x_2 &= p(1-q) - D \\ x_3 &= (1-p)q - D \\ x_4 &= (1-p)(1-q) + D\,. \end{aligned} \tag{23}$$

The mean fitness can be rewritten as

$$\bar{w} = 1 + \beta[(2p-1)(2q-1) + 2D]\,. \tag{24}$$

Straightforward computations (Supplementary *Mathematica* notebook SupplNotebookUnparadox.nb, Section 1.2) yield the following set of equations for the change in allele frequencies and in LD caused by recombination and selection:

$$\bar{w}\Delta^{(rs)}p = \beta p(1-p)(2q-1)\,, \tag{25a}$$

$$\bar{w}\Delta^{(rs)}q = -(\gamma-\beta)q(1-q)(2p-1) + \gamma(2q-1)D\,, \tag{25b}$$

$$\begin{aligned}\bar{w}^2\Delta^{(rs)}D = {} & -(\gamma-\beta)p(1-p)q(1-q) \\ & - \Big\{[1+\beta(2p-1)(2q-1)][\delta+\beta(2p-1)(2q-1)] \\ & \quad + \beta p(1-p)[\gamma - 2(\beta+\gamma)q(1-q)]\Big\}D \\ & + \Big\{\gamma - \beta + \beta(\gamma-3\beta)(2p-1)(2q-1) - 2\beta\delta\Big\}D^2 \\ & + 2\beta(\gamma-\beta)D^3\,. \end{aligned} \tag{25c}$$

The term on the right-hand side of eq. (25a) represents direct selection on locus A. Its strength depends on the frequency of the alleles at locus B, as signified by the factor $(2q-1)$. In eq. (25b), the first additive term on the right-hand side is due to direct selection on locus B, which is again frequency dependent, whereas the second term arises from indirect selection on B transmitted by linkage disequilibrium with A. The equation (25c) for the change of linkage disequilibrium is complicated but shows immediately that if $D = 0$ (and all alleles are present), then $\Delta^{(rs)}D < 0$ if $\gamma > \beta$, and $\Delta^{(rs)}D > 0$ if $\gamma < \beta$. In Section 3.2, we will show a much stronger result.

#### 2.2. Conversion, selection and mutation

With selection, recombination, and mutation, the between-generation changes of $p$, $q$, and $D$ can be written as

$$\Delta p = \Delta^{(rs)}p - 2\mu_A(\Delta^{(rs)}p + p - \tfrac{1}{2})\,, \tag{26a}$$

$$\Delta q = \Delta^{(rs)}q - 2\mu_B(\Delta^{(rs)}q + q - \tfrac{1}{2})\,, \tag{26b}$$

$$\Delta D = \Delta^{(rs)}D - 2[\mu_A(1-\mu_B) + (1-\mu_A)\mu_B](\Delta^{(rs)}D + D)\,. \tag{26c}$$

This is equivalent to eq. (20). If mutation is sufficiently much weaker than selection and recombination, terms of order 2 and higher in the mutation probabilities can be ignored, which further simplifies these equations.

To gain some intuition on the dynamics of this system along the oscillatory orbits, we can assume that linkage disequilibrium is negligible ($D \approx 0$) and rewrite eqs. (26a) and (26b) as follows:

$$\Delta p \approx \tfrac{1}{\bar{w}} bfp(1-p)(1-2\mu_A)(q-\tfrac{1}{2}) - \tfrac{1}{\bar{w}}2\mu_A(p-\tfrac{1}{2})\,, \tag{27a}$$

$$\Delta q \approx -\tfrac{1}{\bar{w}} b(\tfrac{1}{2}c - f)q(1-q)(1-2\mu_B)(p-\tfrac{1}{2}) - \tfrac{1}{\bar{w}}2\mu_B(p-\tfrac{1}{2})\,. \tag{27b}$$

From inspecting these equations it can be concluded that the contribution of fertility $f$ to changes in the frequency of $p$, that is $\Delta p$, is positive when $q > \frac{1}{2}$ and negative otherwise

$$(\Delta p)_f \approx \underbrace{\tfrac{1}{\bar{w}} bp(1-p)(1-2\mu_A)}_{(+)}(q-\tfrac{1}{2})f\,. \tag{28}$$

In the parameter region where oscillations are observed ($f < \frac{1}{2}c$), the contribution of conversion relative to fertility $\frac{1}{2}c - f$ to changes in the frequency of $q$, that is $\Delta q$, is positive when $p < \frac{1}{2}$ and negative otherwise

$$(\Delta q)_c \approx \underbrace{-\tfrac{1}{\bar{w}} bq(1-q)(1-2\mu_B)}_{(-)}(p-\tfrac{1}{2})\underbrace{(\tfrac{1}{2}c - f)}_{+}\,. \tag{29}$$

In addition, the contribution of mutation at the modifier locus $A$ to changes in the frequency of $p$ near the corners of the allelic space, is positive when $p < \frac{1}{2}$ and negative otherwise

$$(\Delta p)_{\mu_A} \approx \underbrace{-\tfrac{1}{\bar{w}}}_{-} 2(p-\tfrac{1}{2})\mu_A\,. \tag{30}$$

Similarly the contribution of mutation in the target locus $B$ to changes in the frequency of $p$, near the corners of the allelic space, is positive when $q < \frac{1}{2}$ and negative otherwise.

$$(\Delta q)_{\mu_B} \approx \underbrace{-\tfrac{1}{\bar{w}}}_{-} 2(q-\tfrac{1}{2})\mu_B\,. \tag{31}$$

Notice that this analysis allows us to develop an intuition regarding the contribution of each parameter to the behaviour of the system but it is far from being rigorous. However, our analysis provides support to the validity of this intuitive approach.

### 3. Deterministic one-target model: results

#### *3.1. Equilibria and their stability*

Much of the subsequent analysis will rely on a perturbation approach that assumes that mutation is much weaker than selection and recombination. In this case, we assume that $\mu_A$ and $\mu_B$ are sufficiently small such that terms of order $\mu_A^2$, $\mu_B^2$, and $\mu_A\mu_B$ can be ignored. Formally, a weak-mutation approximation or a weak-mutation perturbation of an equilibrium is achieved by assuming $\mu_A = \epsilon m_A$ and $\mu_B = \epsilon m_B$, where $m_A \geq 0$ and $m_B \geq 0$ are fixed, and then performing a series expansion of the relevant expressions to first order in $\epsilon$. Finally, the substitutions $m_A \to \mu_A/\epsilon$ and $m_B \to \mu_B/\epsilon$ yield the desired approximation or perturbation in terms of the original parameters.

#### *3.1.1. Corner equilibria*

In the absence of mutation, there exist the four corner equilibria

$$\begin{aligned} \mathbf{x}^{*1} &= (1,0,0,0)\,, \quad \mathbf{x}^{*2} = (0,1,0,0)\,, \\ \mathbf{x}^{*3} &= (0,0,1,0)\,, \quad \mathbf{x}^{*4} = (0,0,0,1)\,, \end{aligned} \tag{32}$$

which correspond to fixation of the gametes $A_1B_1$, $A_1B_2$, $A_2B_1$, and $A_2B_2$, respectively. It was shown in reference (4) that if $\beta > \gamma$, then $\mathbf{x}^{*1}$ and $\mathbf{x}^{*4}$ are linearly stable and $\mathbf{x}^{*2}$ and $\mathbf{x}^{*3}$ are unstable; if $\gamma > \beta$, then all four corner equilibria are saddles. Mutation will perturb these corner equilibria which can either leave the simplex or remain in it.

To determine the fate of the corner equilibria when mutation is added, we perform a straightforward perturbation analysis. As is expected from general theory (e.g., ref. 8, Theorem 4.4), it shows that corner equilibria that are unstable in the absence of mutation leave the simplex if weak mutation is added, and stable corner equilibrium move into the simplex with mutation added. If $\gamma < \beta$, then $\mathbf{x}^{*2}$ and $\mathbf{x}^{*3}$ leave the simplex if mutation is introduced, and the corner equilibria $\mathbf{x}^{*1}$ and $\mathbf{x}^{*4}$ move into the interior of the simplex. If mutation is weak, the coordinates of $\mathbf{x}^{*1}$ are, to leading order in $\mu_A$ and $\mu_B$,

$$x_1^{*1} \approx 1 - [\mu_B\beta + \mu_A(\beta-\gamma)]\,\frac{1+\beta}{\beta(\beta-\gamma)}\,, \tag{33a}$$

$$x_2^{*1} \approx \mu_B\,\frac{1+\beta}{\beta-\gamma}\,, \tag{33b}$$

$$x_3^{*1} \approx \mu_A\,\frac{1+\beta}{\beta}\,, \tag{33c}$$

$$x_4^{*1} \approx 0 \tag{33d}$$

and those of $\mathbf{x}^{*4}$ are

$$x_1^{*4} = x_4^{*1}\,,\; x_2^{*4} = x_3^{*1}\,,\; x_3^{*4} = x_2^{*1}\,,\; x_4^{*4} = x_1^{*1}\,. \tag{34}$$

These two equilibria are linearly stable if mutation is sufficiently weak (8, Theorem 4.4).

If $\gamma > \beta$, then all corner equilibria leave the simplex when mutation is introduced and there are no other equilibria in close vicinity to the corners (8).

*3.1.2. Internal equilibrium*

With or without mutation, there always exists a symmetric internal equilibrium, which we denote by $\mathbf{x}^{*5}$. We start by presenting its coordinates. We note that the probability that no mutation occurs is $(1-2\mu_A)(1-2\mu_B)$ and define the total mutation probability by

$$\mu_{\text{tot}} = 1 - (1-2\mu_A)(1-2\mu_B)\,. \tag{35}$$

Then the symmetric internal equilibrium is given by (Supplementary *Mathematica* notebook SupplNotebookUnparadox.nb, Section 2.5)

$$p^* = q^* = \tfrac{1}{2} \tag{36a}$$

and

$$D^* = \frac{2\delta(1-\mu_{\text{tot}}) + 2\mu_{\text{tot}} - \sqrt{R}}{4(\gamma-\beta) - 4\mu_{\text{tot}}(\beta+\gamma)}\,, \tag{36b}$$

where

$$R = 4[\delta(1-\mu_{\text{tot}}) + \mu_{\text{tot}}]^2 + \gamma^2(1-\mu_{\text{tot}})^2 - 2\beta\gamma(1-\mu_{\text{tot}}) + \beta^2(1-\mu_{\text{tot}}^2)\,. \tag{36c}$$

It is straightforward to show that $D^* < 0$ if and only if $\gamma > \beta$ (also in the exceptional case $\mu_{\text{tot}} = \dfrac{\gamma-\beta}{\beta+\gamma}$), $D^* > 0$ if and only if $\gamma < \beta$, and $D^* = 0$ if and only if $\gamma = \beta$.

If $\mu_{\text{tot}} = \dfrac{\gamma-\beta}{\beta+\gamma}$ then

$$D^* = -\frac{\beta(\gamma-\beta)}{8(\gamma-\beta+2\beta\delta)}\,, \tag{36d}$$

which is also negative because this case requires $\gamma > \beta$.

Assuming weak mutation, we observe that $\mu_{\text{tot}} \approx 2(\mu_A+\mu_B)$. A simple perturbation analysis, as outlined above, yields the following first-order approximation for $D^*$:

$$D^* \approx D_0^* + (\mu_A+\mu_B)\,D_1^*\,, \tag{37a}$$

where

$$D_0^* = -\frac{\sqrt{(\gamma-\beta)^2+4\delta^2}-2\delta}{4(\gamma-\beta)} \tag{37b}$$

and

$$D_1^* = \frac{2(\gamma-\beta+2\beta\delta)\left(\sqrt{(\gamma-\beta)^2+4\delta^2}-2\delta\right)-\beta(\gamma-\beta)^2}{2(\gamma-\beta)^2\sqrt{(\gamma-\beta)^2+4\delta^2}}\,. \tag{37c}$$

We note that the leading term $D_0^*$, i.e., the linkage disequilibrium at the internal equilibrium in the absence of mutation, is negative if and only if $\gamma > \beta$ (cf. ref. 4), and the coefficient $D_1^*$ of $\mu_A+\mu_B$ is positive if and only if $\gamma > \beta$ (provided $\gamma \le 1$). Therefore, mutation weakens linkage disequilibrium by reducing its absolute value.

Under a weak perturbation, an internal equilibrium maintains its stability properties if the unperturbed equilibrium is hyperbolic (e.g., 8, Theorem 4.4). Hyperbolic means that there is no eigenvalue of modulus 1. Because in the absence of mutation, the symmetric internal equilibrium is linearly stable if $\gamma > \beta$ (4), its perturbation given in eq. (36) remains linearly stable. If $\gamma < \beta$, then it remains a saddle point under weak mutation (and all eigenvalues are real). Furthermore, in the absence of mutation $\mathbf{x}^{*5}$ is the only internal equilibrium, i.e., the only equilibrium at which all four gametes are present.

If $\gamma > \beta$, then the condition

$$\delta > \frac{(\gamma - 2\beta)^2}{8\gamma}\sqrt{\frac{\gamma - \beta}{\beta}} \tag{38}$$

implies that, in the absence of mutation, the symmetric equilibrium has a pair of conjugate complex (i.e., non-real) eigenvalues. Hence, the symmetric equilibrium is a spiral sink in this case. This property is maintained under weak perturbations because eigenvalues change continuously. Notice that condition (38) is missing in reference (4).

In summary, these results shows that if $\gamma > \beta$, which is equivalent to $f < \frac{1}{2}c$, then there is no equilibrium close to the boundary and the symmetric internal equilibrium $\mathbf{x}^{*5}$ is linearly stable. In addition, solutions starting close enough to it spiral towards it if condition (38) is satisfied, which is the case if the 'effective' recombination probability $\delta$ is not too small. Because in the absence of mutation, a stable heteroclinic orbit can exist if $\gamma > \beta$ (4), with mutation it may be replaced by a stable limit cycle close to the boundary. We have not proved this, but numerical work suggests existence and (local) stability of a limit cycle (see Figures S2 and 4 in the Main Text). If $\gamma < \beta$, the symmetric internal equilibrium is unstable, in fact a saddle point, and two equilibria close to fixation of the gametes $A_1B_1$ and $A_2B_2$ exist and are linearly stable. Numerical work suggests that every solution converges to one of these nearly monomorphic equilibria. Next we show that any attractor exhibits negative linkage disequilibrium.

#### *3.2. Attractors exhibit negative linkage disequilibrium if $\gamma > \beta$ and $\delta \geq \beta$*

Throughout this subsection, we assume $\gamma > \beta$ and $\delta \geq \beta$. As a first step, we prove that $D \geq 0$ implies $\Delta^{(rs)}D \leq 0$ and $\Delta^{(rs)}D = 0$ only on the edges at which one locus is fixed.

Let $D \geq 0$. The right-hand side of (25c) can be written as $d_1 + d_2 + d_3 + d_4$, where

$$\begin{aligned}
d_1 &= -(\gamma - \beta)p(1-p)q(1-q) + (\gamma - \beta)D^2\,, \\
d_2 &= -[1 + \beta(2p-1)(2q-1)][\delta + \beta(2p-1)(2q-1)]D\,, \\
d_3 &= -\beta p(1-p)[\gamma - 2(\beta+\gamma)q(1-q)]D + 2\beta(\gamma-\beta)D^3\,, \\
d_4 &= [\beta(\gamma - 3\beta)(2p-1)(2q-1) - 2\beta\delta]D^2\,.
\end{aligned}$$

Because $D^2 \leq p(1-p)q(1-q)$ and $\gamma > \beta$, we have $d_1 \leq 0$ and $d_1 = 0$ only if one of the allele frequencies vanishes. Because $\beta < \min\{\delta, 1\}$, each of the two factors in brackets is positive and we conclude that $d_2 \leq 0$ if $D \geq 0$. Next, we rewrite $d_3$ as

$$d_3 = -\beta D \left\{\gamma p(1-p) - 2(\beta+\gamma)p(1-p)q(1-q) - 2(\gamma-\beta)D^2\right\}\,.$$

Then, because $D^2 \leq p(1-p)q(1-q)$, we derive

$$\begin{aligned}
&\gamma p(1-p) - 2(\beta+\gamma)p(1-p)q(1-q) - 2(\gamma-\beta)D^2 \\
&\qquad \geq \gamma p(1-p) - 2p(1-p)q(1-q)[(\beta+\gamma) + (\gamma-\beta)] \\
&\qquad = \gamma p(1-p)[1 - 4q(1-q)] \geq 0.
\end{aligned}$$

Therefore, $d_3 \leq 0$. Finally, we have $\beta(\gamma - 3\beta)(2p-1)(2q-1) - 2\beta\delta \leq \beta(|\gamma - 3\beta| - 2\delta)$. If $\gamma \leq 3\beta$, then $|\gamma - 3\beta| - 2\delta = 3\beta - \gamma - 2\delta = 2(\beta - \delta) + \beta - \gamma < 0$. If $\gamma > 3\beta$, then $|\gamma - 3\beta| - 2\delta = \gamma - 3\beta - 2\delta < 0$ by using $\delta \geq \frac{1}{2}\gamma$ in (10). This shows that $d_4 \leq 0$. Therefore, $\Delta^{(rs)}D \leq 0$ if $D \geq 0$ and $\Delta^{(rs)}D < 0$ unless at least one of the alleles is absent.

If mutation is weak (but present), so that the higher-order mutation terms in (26c) can be ignored, it follows immediately that $\Delta D < 0$ if $D \geq 0$. Therefore, the statement that attractors exhibit negative linkage disequilibrium follows.

The average population mean fitness in a complete cycle is always greater than the population mean fitness corresponding to the internal equilibrium (Figure S3).

### 4. Deterministic two-target (three-locus) model

In this section, we consider a three-locus model that extends the previous two-locus model by adding a second locus (locus $C$) as potential target of the PRDM9-like protein encoded in locus $A$. We assume that during meiosis the PRDM9-like protein attempts to bind one and only one target allele either at locus $B$ or $C$ with equal probability $\frac{1}{2}$. As a result, in our model there is at most one crossover per meiosis. We assume that the interaction between locus $A$ and $C$ is identical to the one between $A$ and $B$. Binding between the protein encoded by locus $A$ and target $C$ happens with probability $b_{i,m}$. Binding between protein and target leads to a double-strand break, which may result in crossover between the flanking regions of the target locus with probability $r_t$ and conversion with probability $c$. The rest of the life cycle is the same as in the two-locus model.

Hereafter we assume that a match between subscripts of alleles at locus $A$ and $C$ is required for binding between protein and target and double-strand break at the target with probability $b$, that is:

$$b_{i,m} = \begin{cases} b & \text{if } i = m \\ 0 & \text{if } i \neq m. \end{cases}$$

Due to the increased complexity of the three-locus model, for clarity we separate the steps of recombination between loci ($A$,$B$,$C$) and crossover in each of the target ($B$ and $C$) between their flanking regions.

*4.1. Crossover*

Let $r_{(A,B)}$, $r_{(B,C)}$, and $r_{(A,C)}$ be the probabilities of recombination between loci $A$ and $B$, $B$ and $C$, $A$ and $C$, respectively. As in previous sections we assume that the loci are either far away from each other on the same chromosome or on separate chromosomes. Therefore we assume $r_{(A,B)} = r_{(B,C)} = r_{(A,C)} = r_m \approx \frac{1}{2}$.

The frequency of genotype $\frac{A_i B_k C_m}{A_j B_l C_n}$ after recombination is

$$\begin{aligned} (x_{i,k,m} x_{j,l,n})^{(r)} &= (1 - r_m)^2 x_{i,k,m} x_{j,l,n} \\ &\quad + r_m (1 - r_m)(x_{i,l,n} x_{j,k,m} + x_{i,k,n} x_{j,l,m}) + r_m^2 x_{i,l,m} x_{j,k,n} \end{aligned} \tag{39}$$

*4.2. Conversion and selection*

During meiosis, double-strand breaks may occur in targets $B$ and $C$, which can trigger conversion and influence proper chromosomal segregation.

The frequency of the gametic type $A_i B_k C_m$ after meiosis is given by

$$\begin{aligned} x_{i,k,m}^{(rs)} = \tfrac{1}{\bar{w}} \textstyle\sum_{j,l,n} & \Big[\Big((1-c)\,\bar{\bar{b}}_{ij,kl} + (1-f)\,(1 - \bar{\bar{b}}_{ij,kl}) \\ & + \tfrac{1}{2} c \left(\bar{b}_{ij,l} + \bar{b}_{ij,n}\right)\Big) \left(x_{i,k,m} x_{j,l,n}\right)^{(r)} \\ & + \tfrac{1}{2} c \Big(\bar{b}_{ij,l} \left(x_{i,l,m} x_{j,k,n}\right)^{(r)} + \bar{b}_{ij,n} \left(x_{i,k,n} x_{j,l,m}\right)^{(r)}\Big)\Big] \end{aligned} \tag{40}$$

where

$$\bar{w} = \sum_{i,k,m} \sum_{j,l,n} \left[\bar{\bar{b}}_{ij,kl} + (1-f)\,(1 - \bar{\bar{b}}_{ij,kl})\right] \left(x_{i,k,m} x_{j,l,n}\right)^{(r)} \tag{41}$$

*4.3. Mutation*

Following previous definitions we denote by $\mu_{C,mn}$ the mutation probability from $C_m \to C_n$ where $m \neq n$. For simplicity, we assume that mutations occur as frequently in one direction as in the other, that is $\mu_{A,ij} = \mu_{A,ji} = \mu_A$, $\mu_{B,kl} = \mu_{B,lk} = \mu_B$ and

$\mu_{C,mn} = \mu_{C,nm} = \mu_C$. Finally, we assume that mutations are sufficiently weak to ignore multiple mutations.

The frequency of the gametic type $A_iB_kC_m$ at the beginning of the next generation is given by

$$\begin{aligned} x'_{i,k,m} &= x^{(rs)}_{i,k,m}\left(1 - \sum_{j:j\neq i}\mu_A - \sum_{l:l\neq k}\mu_B - \sum_{n:n\neq m}\mu_C\right) \\ &\quad + \sum_{j:j\neq i} x^{(rs)}_{j,k,m}\mu_A + \sum_{l:l\neq k} x^{(rs)}_{i,l,m}\mu_B + \sum_{n:n\neq m} x^{(rs)}_{i,k,n}\mu_C\,. \end{aligned} \tag{42}$$

These changes in frequency over the whole life cycle underpin changes in the population mean crossover at targets $B$ and $C$ (target crossover), referred to as $\overline{r}_{t(B)}$ and $\overline{r}_{t(C)}$ respectively, and the population mean crossover across all targets (genome crossover), referred to as $\overline{r}_t$. The expression for these phenotypes are

$$\overline{r}_{t(B)} = \sum_{i,k,m}\sum_{j,l,n} \overline{\overline{b}}_{ij,kl}\ r_t\ x_{i,k,m}\ x_{j,l,n} \tag{43a}$$

$$\overline{r}_{t(C)} = \sum_{i,k,m}\sum_{j,l,n} \overline{\overline{b}}_{ij,mn}\ r_t\ x_{i,k,m}\ x_{j,l,n} \tag{43b}$$

$$\overline{r}_t = \tfrac{1}{2}\overline{r}_{t(B)} + \tfrac{1}{2}\overline{r}_{t(C)} \tag{43c}$$

### 5. Stochastic multi-target model

To investigate how the system behaves with multiple targets and in finite populations, we built a stochastic version of the deterministic models discussed. We assume that, PRDM9 and all target loci can have 2 alleles, $A_1$ and $A_2$ for PRDM9, $B_1$ and $B_2$ for all the targets. We still assume that there is only one binding attempt per meiotic event, and that a binding may be successful if and only if the subscripts of the alleles in locus A and B match.

We consider a population of $N$ individuals. This population is initialized with frequencies $q_0^2$ of $B_1B_1$, $q_0(1-q_0)$ of $B_1B_2$, $q_0(1-q_0)$ of $B_2B_1$, and $(1-q_0)^2$ of $B_2B_2$ at each target locus. Also, each PRDM9 allele in the initial population has probability $p_0$

of being $A_1$ and $1 - p_0$ of being $A_2$. In all simulations presented here, we set $q_0 = 0.95$ and $p_0 = 0.99$. Consequently, the simulations are always initialized with all targets in hot state.

The life cycle of the population is the same as in the deterministic model. During meiosis, recombination occurs, shuffling genetic associations. We assume a recombination rate of $\frac{1}{2}$ between each pair of loci (that is all loci recombine freely) unless specified otherwise.

After recombination, a target locus is chosen uniformly at random to be the actual target of a binding attempt. One of the two allelic copies present in the PRDM9 locus is chosen with probability $\frac{1}{2}$ of being the copy that is attempting to bind a target, and one of the two allelic copies present in the target locus is chosen with probability $\frac{1}{2}$ of being the copy that experiences a binding attempt. If the two chosen alleles match, the binding is successful with probability $b$. In all simulations presented here, $b$ is set to 1. In the case of a successful attempt, with probability $c$ the bound target allele is converted to its homologous copy, and the fitness of the individual is set at 1. In contrast, if binding is not successful, there is no conversion and the fitness of the individual is set at $1-f$. In all simulations presented here we assume that $f = 0.4$.

After recombination and conversion, segregation is modelled by randomly drawing one of the two possible types of meiotic product. Pending mutation, this is the gamete used to produce offspring.

Mutation in these gametes occurs with probability $\mu_A$ on the PRDM9 locus, and with probability $\mu_B$ on each target locus. A mutation always transforms one allele into the other.

Selection is modelled when determining the parents of an individual of the next-generation population. Each parent is chosen in a current population by drawing it from a uniform distribution. Once it is chosen, we run its meiosis and calculate its fitness. It is then retained with a probability equal to its fitness; if discarded, we choose a new parent in the same way, and iterate this process of choosing and discarding until a parent is retained. To produce an individual, two parents must be retained. The selection procedure is iterated until $N$ parents have been produced.

This life cycle is iterated during a large number of generations. We use the first 2,000 generations as a burn-in phase, during which we assume no actual effect of PRDM9 and target alleles: $f = 0$ and $c = 0$ during this phase. This allows to shuffle the whole population and reach a mutation-recombination-drift equilibrium. After the 2,000 burn-in generations, $f$ and $c$ are set at their values, and we let the system evolve.

Every 10 generations, we store in separate files: (1) for each target locus the probability of a successful PRDM9–target binding, (2) the frequency of the $A_1$ allele, (3) the frequency of all $B_1$ alleles. (1) is calculated by checking the genomes of all individuals in the population, and assuming that there is a probability $\frac{b}{4N_{\mathrm{loc}}}$ of a successful binding each time a PRDM9 allele matches a target allele. Here, $N_{\mathrm{loc}}$ designates the number of target loci considered. This comes from the fact that one given PRDM9 allele has a probability $\frac{1}{4N_{\mathrm{loc}}}$ of attempting to bind a particular target allele, and a probability of $b$ of succeeding if the alleles match.

In Fig. S5, we present the periods of the oscillations of the crossover probability (which equals the probability of a successful binding (1)) at each target locus. Because here population size is finite, so that allele frequencies and crossover probabilities vary stochastically, we calculated the periods of the oscillations in a different way than for the deterministic model. The algorithm finds the first generation of a shift from a hot (high crossover probability) to a cold state (low crossover probability), and then calculates the number of generations between each of the following shifts. The mean and variance of these numbers of generations between shift are shown in Fig. S5 as the mean and variance of the period of the oscillations. Figure S5 illustrates that the stochasticity of the model induces ample variation in the number of generations between two successive shifts from hot to cold. To ensure that the variances are calculated over a similar number of cycles, we performed the computations for panel (i) in Fig. S5 for 10 000 generations if $\mu_A = \mu_B = 10^{-6}$, for 100 000 generations if $\mu_A = \mu_B = 10^{-7}$, and for 1 000 000 generations if $\mu_A = \mu_B = 10^{-8}$.

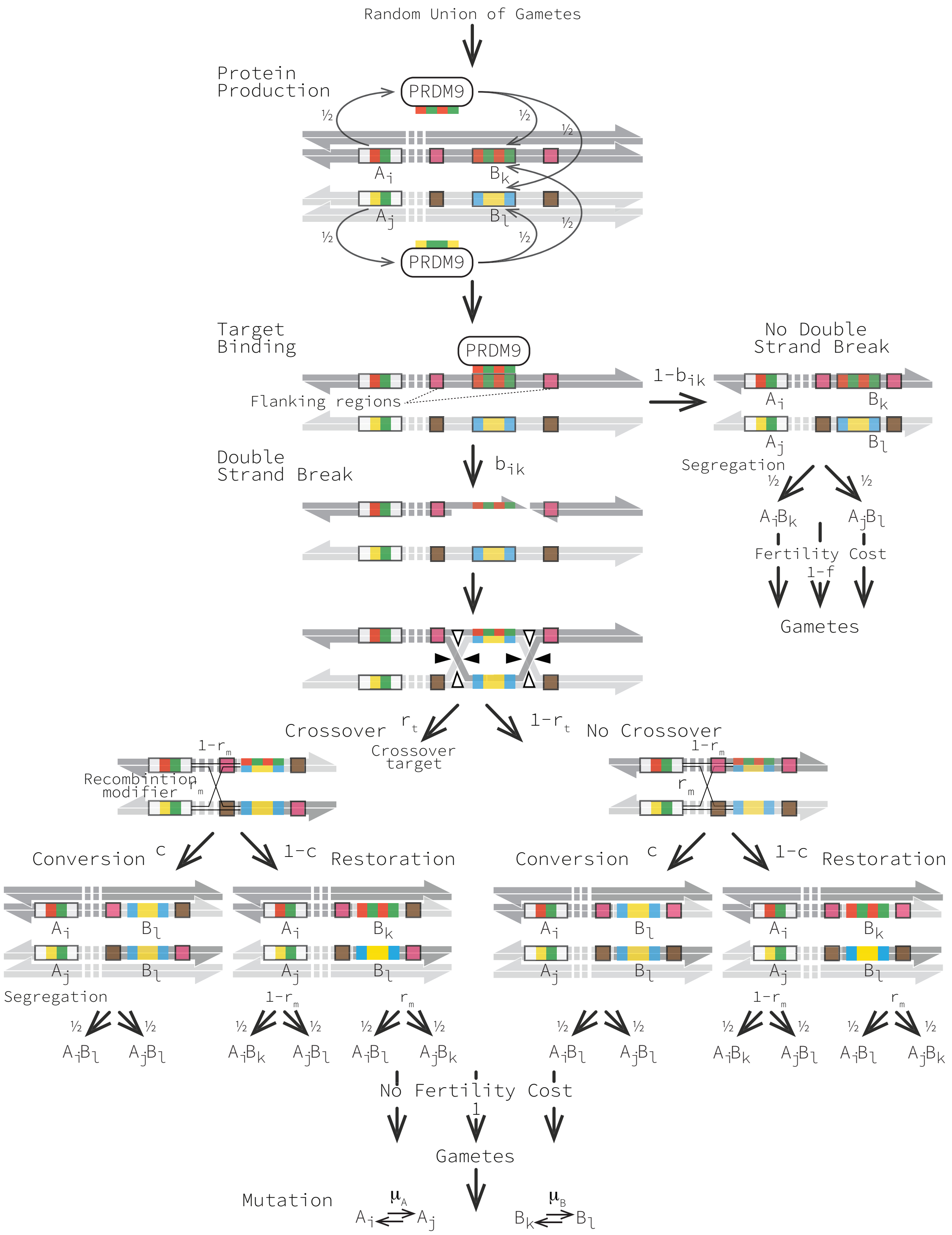


Figure S1: Model of recombination initiated by a double strand break (DSB) in a specific target sequence. The life cycle starts with the production of a PRDM9-like protein with a recognition sequence that may produce a double-strand break (DSB) when matching the target motif (represented by the same colour of sequence and motif) with probability $b_{ik} = b$ in the main text. If protein and target bind, our model follows the canonical DSB repair model for the initiation of recombination (5, 6). Once recombination has been completed, alleles segregate into gametes following Mendel's laws. If there is no recombination alleles may not segregate properly into gametes. Finally mutations take place. Notice that sister chromatids are represented in the first and last step but are omitted in intermediate steps for clarity.

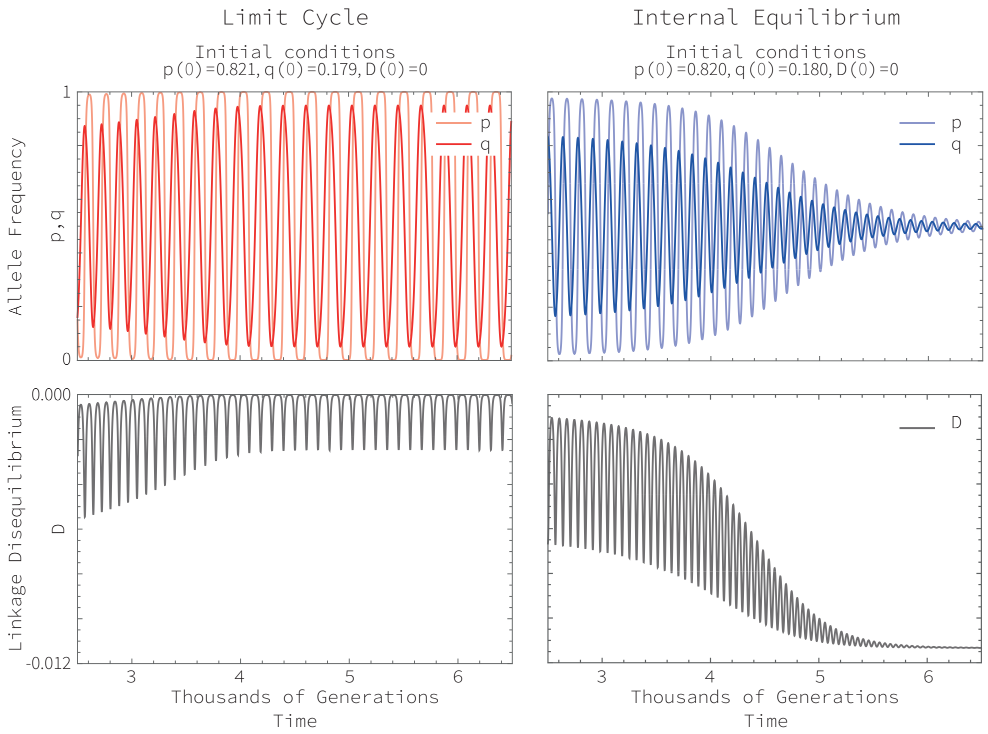


Figure S2: Examples of time series of the allelic frequencies in the parameter region $f < c/2$. The first column corresponds to initial conditions leading to a stable limit cycle. In the second column the initial conditions are slightly different leading to an internal equilibrium. The first row shows the frequency of alleles $A_1$ ($p$) and $B_1$ ($q$). The second row shows the linkage disequilibrium between the different gametes ($D$). If the allele frequencies converge to a cycle, the linkage disequilibrium also does. In this cycle the linkage disequilibrium is always negative but very small. If the allele frequencies converge to the internal equilibrium, the linkage disequilibrium tends to a small negative value. The trajectories correspond to the set of parameter values $f = 0.4$, $b = c = 1$, $r_m = \frac{1}{2}$, $\mu = 10^{-5}$.

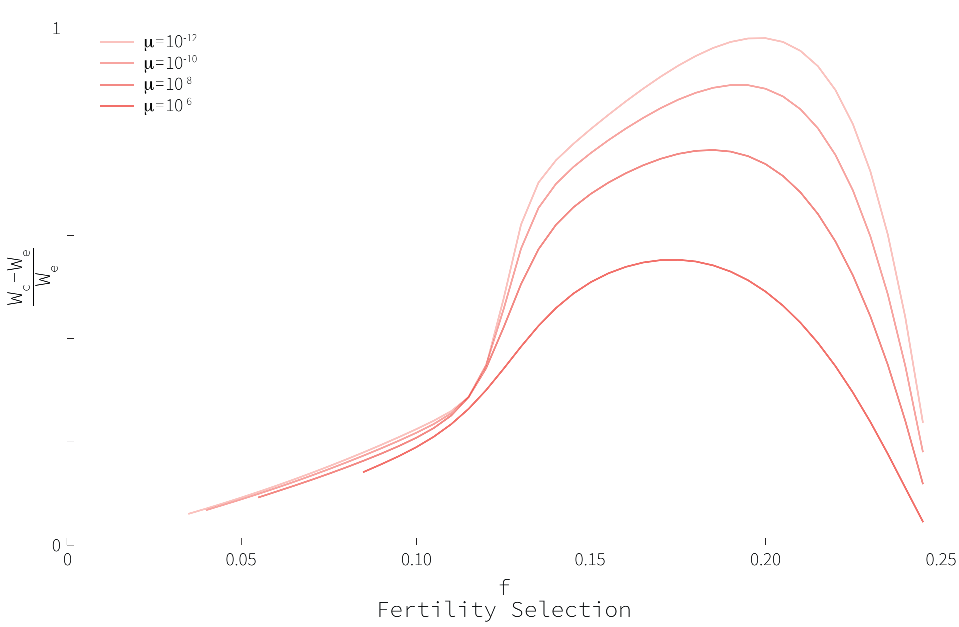


Figure S3: Relative difference between the average of the population mean fitness during a cycle and the population mean fitness at the internal equilibrium.

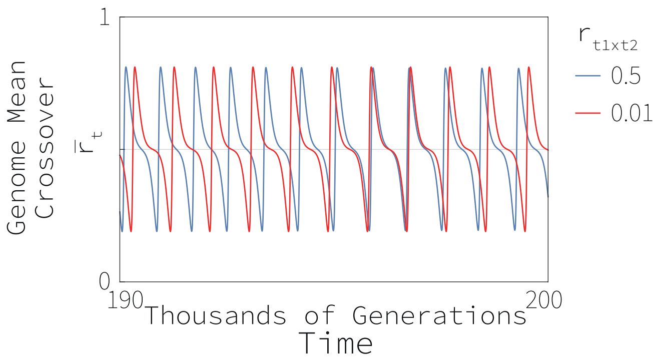


Figure S4: Effect of the recombination probability between targets on the genomic mean crossover rate for the deterministic one-modifier two-targets model with two alleles at each locus. The blue curve corresponds to the maximum crossover probability of $\frac{1}{2}$ ($r_{t1\times t2} = 0.5$), the red line to a small crossover probability of 0.01 ($r_{t1\times t2} = 0.01$). Except for slightly faster oscillations when the crossover probability is reduced, there is no qualitative difference between the dynamics. The parameter values used to create this figure are $f = 0.2$, $b = 1$, $c = \frac{1}{2}$, $r_m = \frac{1}{2}$, $\mu = 10^{-8}$. Initial frequencies of haplotypes are: $x_{1,1,1}(0) = 0.425$, $x_{1,1,2}(0) = 0.238$, $x_{1,2,1}(0) = 0.208$, $x_{1,2,2}(0) = 0.021$, $x_{2,1,1}(0) = 0.033$, $x_{1,1,2}(0) = 0.021$, $x_{1,2,1}(0) = 0.033$, $x_{1,2,2}(0) = 0.021$.

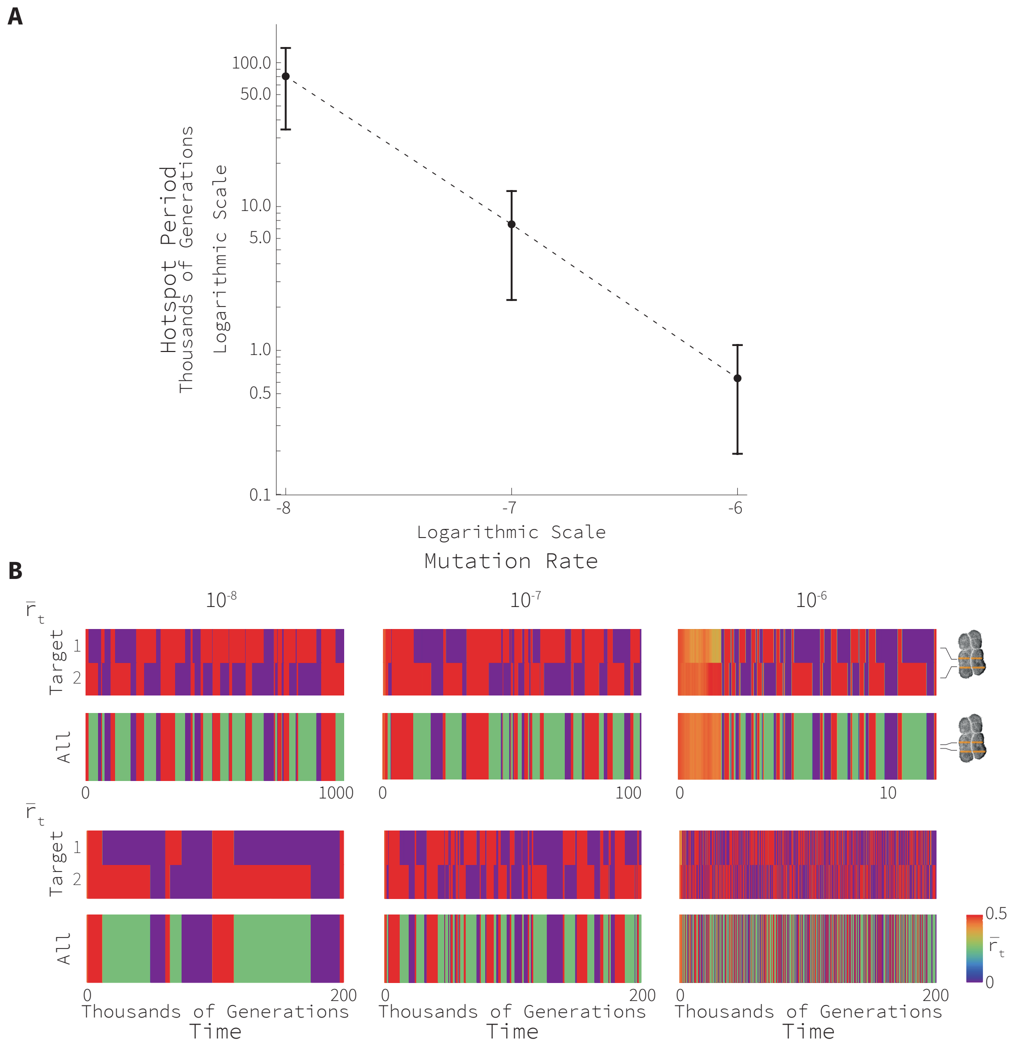


Figure S5: Effect of the mutation probability on the rate of hotspot oscillations in a finite population. Panel (A) shows the average period of an oscillation between hot and cold states for different mutation probabilities. Dots represent the average value of the period (computed for 10 000 generations if $\mu_A = \mu_B = 10^{-6}$, for 100 000 generations if $\mu_A = \mu_B = 10^{-7}$, and for 1 000 000 generations if $\mu_A = \mu_B = 10^{-8}$) for a population size of 5 000. Vertical bars show the standard deviation. Panel (B) shows the change in recombination over time in each target and the average for the genome. We explored three mutation rates that differ in one order of magnitude, namely $10^{-8}$, $10^{-7}$, and $10^{-6}$. The first row shows the hotspots dynamics for the periods of time 1 000 000, 100 000, and 10 000 generations; time changes one order of magnitude with their corresponding mutation probability. This row illustrates how increasing the sampling time one order of magnitude produces similar number of oscillation in the three mutation probabilities considered. In the second row we show the dynamics when the sampling time remains constant. This row illustrates how mutation probabilities increase the rate of oscillations. The parameter values used to create this figure are $f = 0.4$, $b = c = 1$, $r_m = \frac{1}{2}$, and $p(0) = 0.95$, $q(0) = 0.99$, $D(0) = 0$ and $N = 5,000$
